# Bile acid signaling as a therapeutically tractable pathway linking early caregiving adversity to social behavior

**DOI:** 10.64898/2026.08.14.742173

**Authors:** Cesar A. Medina, Pragney Deme, Vishal Win, Isabellah Nikitah, Isaac McKie, Mingyuan Song, Maria Raudales, Elaina Regier, Emily Amsden, Pranali Bendale, Norman Haughey, Maya Opendak

**Affiliations:** Solomon H. Snyder Department of Neuroscience, The Johns Hopkins University School of Medicine, Baltimore, MD, USA 21205; Kennedy Krieger Institute, Baltimore, MD, USA 21205; Department of Physiology, Tulane University School of Medicine, New Orleans, LA, USA 70112; Department of Neurology, The Johns Hopkins University School of Medicine, Baltimore, MD, USA 21205; Department of Psychiatry, The Johns Hopkins University School of Medicine, Baltimore, MD, USA 21205; The Brain Institute, Tulane University School of Medicine, New Orleans, LA, USA 70112; Kavli NeuroDiscovery Institute, Baltimore MD USA 21218

**Author notes:** Corresponding Author: Maya Opendak PhD, 707 N. Broadway, Rm. 400P, Baltimore MD USA 21205,.

## Abstract

Adverse early caregiving produces lasting changes in social behavior and increases vulnerability to psychiatric illness, yet the biological pathways through which these experiences become embedded during development remain poorly understood. Using two complementary rat models of early-life adversity (ELA), we combined behavioral phenotyping with serum metabolomics and basolateral amygdala (BLA) transcriptomics to identify bile acid biology as a candidate pathway associated with disrupted social development. In the Deconstructed Adversity Model (DAM), which dissociates adverse social experience from non-social stress, social adversity produced distinct behavioral alterations accompanied by sex-, age-, and adversity-dependent changes in peripheral bile acid and tryptophan metabolism together with sex-specific BLA gene co-expression networks associated with social behavior. These coordinated peripheral and central alterations converged on bile acid biology as a candidate pathway for pharmacological intervention. We next tested this prediction in the Scarcity Adversity Model via Limited Bedding (SAM-LB), where oral supplementation with chenodeoxycholic acid (CDCA), but not the related primary bile acid cholic acid (CA), during the adversity period rescued the infant affiliative social deficit produced by adverse caregiving. Together, these findings identify bile acid biology as a pharmacologically tractable pathway associated with the developmental consequences of adverse early caregiving and support further investigation of CDCA, an FDA-approved bile acid, as a candidate intervention for mitigating early social behavioral deficits.

## INTRODUCTION

Early social experience is a powerful organizer of brain development.^1,2^ Disrupted caregiving increases the lifelong risk of anxiety, depression, and other neuropsychiatric disorders characterized by impaired social behavior.^3,4^ Across species, adverse early caregiving alters the developmental trajectory of social circuits, producing lasting changes in how individuals respond to caregivers, peers, and social threat.^5–8^ Rodent models have been instrumental in identifying the neural circuits through which early-life adversity (ELA) shapes later behavior,^9–11^ yet the peripheral biological pathways through which adverse caregiving becomes biologically embedded remain poorly understood. Peripheral metabolic signals ultimately shape neural circuit function through changes in neuronal signaling and gene expression,^12^ making it essential to understand how adversity reorganizes both peripheral metabolism and central transcriptional programs to produce lasting behavioral change. These biological changes arise during sensitive developmental periods, long before psychiatric symptoms become apparent,^13^ creating an opportunity to identify modifiable peripheral pathways for preventive intervention. Unlike neural circuits, peripheral metabolic pathways may offer therapeutically accessible targets during these windows of developmental plasticity.

A growing body of evidence identifies bile acid (BA) signaling as a compelling candidate pathway linking peripheral metabolism to brain function. Primary bile acids, including cholic acid (CA) and chenodeoxycholic acid (CDCA), are cholesterol-derived endocrine signaling molecules synthesized in the liver that regulate peripheral metabolism while also influencing central nervous system function. Dysregulation of BA metabolism has been implicated in psychiatric and metabolic disorders, including anxiety and depression.^14,15^ In patients with major depressive disorder, circulating CDCA, but not CA, is reduced relative to healthy controls, and BA profiles normalize during remission in parallel with symptom improvement.^14,16–19^ Bile acids influence the central nervous system through multiple signaling mechanisms and regulate neuronal function and gene expression.^20,21^ Collectively, these findings suggest that BA signaling is well positioned to coordinate peripheral metabolic state with central molecular programs relevant to social behavior. However, whether BA signaling contributes to the developmental consequences of adverse early caregiving, or represents a therapeutically targetable pathway for preventing social behavioral deficits, remains unknown.

Evidence from both adult stress models and models of early-life adversity further implicates bile acid signaling in behavioral regulation. In adult rodents, chronic unpredictable mild stress alters bile acid metabolism, and normalization of this pathway accompanies antidepressant treatment, with CDCA emerging as a candidate bioactive metabolite.^22^ Likewise, direct administration of CDCA reduces depression-like behavior,^14^ while social defeat stress produces parallel changes in bile acid composition and social behavior.^23^ Early-life adversity similarly disrupts related physiology. Scarcity Adversity via Limited Bedding (SAM-LB) alters gut microbial composition,^24^ maternal separation elevates circulating bile acids,^25^ and perinatal stress perturbs bile acid metabolism before the emergence of behavioral abnormalities.^26^ Finally, bile acid metabolism is sexually dimorphic and regulated in part by gonadal hormones,^27^ raising the possibility that BA signaling contributes to the well-established sex differences in vulnerability to early-life adversity. Together, these observations suggest that BA signaling is responsive to both stress and adverse caregiving and may represent a biologically meaningful interface between early experience, peripheral metabolism, and behavioral development. Whether this pathway contributes causally to the developmental consequences of adverse caregiving, whether its effects depend on the social context of adversity, and whether it can be therapeutically targeted remain unknown.

To address these questions, we combined two complementary rat models of early-life adversity with distinct experimental strengths. The Deconstructed Adversity Model (DAM) dissociates adverse social experience from non-social stress by comparing repeated shock delivered either alone or in the presence of a maternal odor-producing dam, enabling mechanistic dissection of how adverse social context reshapes peripheral metabolism, basolateral amygdala (BLA) transcriptional programs, and social behavior.^5^ We complemented this approach with the widely used Scarcity Adversity Model via Limited Bedding (SAM-LB), a robust model of adverse caregiving that is well suited for pharmacological intervention studies.^6,9,28–31^ We hypothesized that adverse social experience reorganizes coordinated peripheral metabolic and central transcriptional programs and that pharmacologic modulation of bile acid signaling would modify the resulting behavioral phenotype. Here, we show that adverse social experience is associated with coordinated changes in social behavior, peripheral metabolism, and BLA transcriptional organization and that oral supplementation with chenodeoxycholic acid (CDCA), but not the related primary bile acid cholic acid (CA), rescues the resulting infant social behavioral deficit. Together, these findings identify bile acid signaling as a pharmacologically tractable pathway linking adverse early caregiving to disrupted social behavior.

## RESULTS

### Adverse social experience selectively disrupts affiliative social behavior across development

To determine whether the social context of early adversity, rather than adversity itself, drives long-term alterations in social behavior, we used the Deconstructed Adversity Model (DAM), in which pups experienced repeated shock either alone (non-social adversity, SA), in the presence of a maternal odor-producing dam (social adversity, SM), or a no-shock control condition (beaker alone, BA) from P8–P12 (Fig. 1A).

**Figure 1.**
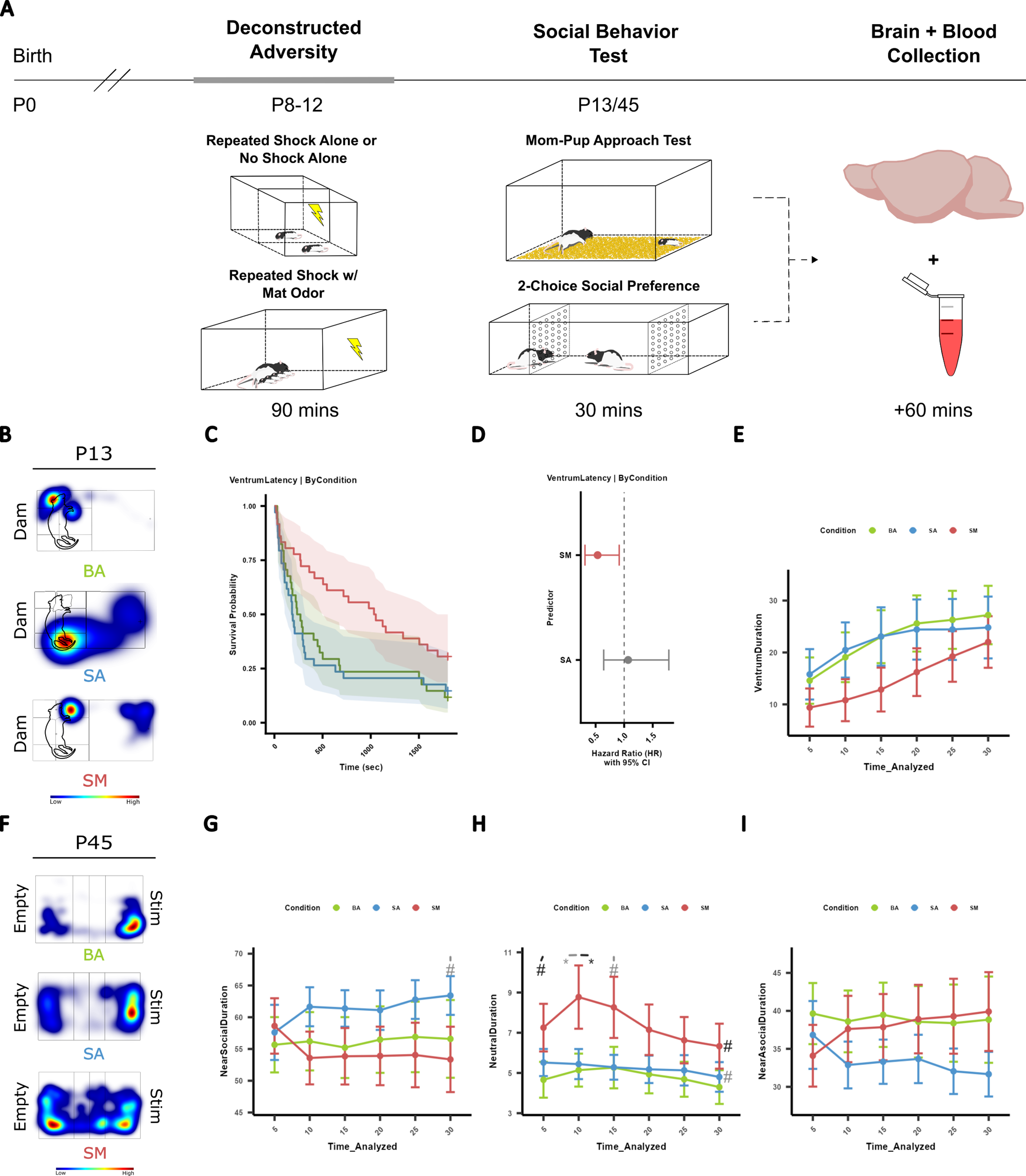
Early social but not non-social stress perturbs infant and adolescent social behavior. (A) Experimental design of behavior and tissue collection following deconstructed adversity model (DAM). Briefly, male and female pups undergo repeated no shock alone (beaker alone, BA), repeated shock alone (shock alone, SA), or repeated shock with maternal odor (i.e., dam, shock mom, SM) treatment for 90 mins each day from P8-P12. On P13, infant social behavior is tested using the maternal social approach test. Brain and blood from half of the pups are randomly collected. On P45, adolescent social behavior is tested using 2-choice social preference test before brain and blood collection. (B) Representative heatmaps of BA, SA, and SM treated infant pups in maternal social approach test show relative position of pups to an anesthetized maternal-odor producing Dam. Dam outline overlaid onto heatmap. Blue and red indicate low and high proportion respectively. (C) Kaplan-Meier survival curves of BA, SA, and SM treated infant pups’ latency to approach Dam. Survival probability (y-axis) of approaching the ventrum is graphed for each treatment as a function of Time (sec). (D) Cox-proportional hazard and 95% confidence interval (CI) of latency to approach the ventrum by SA and SM treated relative to BA treated infant pup. Red indicates significantly different and gray indicates no significant difference. (E) Line graph of proportion of total duration (%) spent around the ventrum by the BA, SA, and SM treated infant pups is plotted for each 5 min time bin. (F) Representative heatmaps of duration near empty or social stim (Stim) side of 2-CSP chamber by BA, SA, and SM treated adolescent pups in 2-CSP. Blue and red indicate low and high proportion respectively. (G, H, I) Line graph of proportion of total duration (%) spent near the social stim (NearSocial), neutral (middle), and near the empty (Empty) side of the 2-CSP apparatus are plotted within each 5 min time bin. Area under the curve of proportion of total duration spent around or within a region of interest is compared across each comparison (BA vs SA, BA vs SM, SM vs SA) and plotted to the right of the 30 min time bin value. Green is BA, blue is SA, and red is SM. # indicates p < 0.1, * indicates p < 0.05, ** indicates p < 0.01, and *** indicates p < 0.001. Additionally, light gray * or # reflect comparison between BA and SA, black * or # reflect comparison between BA and SM, and dark gray * or # reflect comparison between SA and SM.

To assess immediate behavioral consequences, pups underwent a maternal social approach (MSA) test at P13. Consistent with previous use of this measure,^5,30,32^ a subset of animals never approached the dam during the 30-minute session; therefore, latency to approach was analyzed using Kaplan–Meier survival curves and Cox proportional hazards models. Social adversity preferentially disrupted approach to the ventrum, the affiliative region where nursing and grooming occur. SM pups exhibited significantly delayed ventrum approach relative to BA controls (HR = 0.53, 95% CI 0.31–0.91, p = 0.022; global log-rank p = 0.020; Fig. 1B-D; Table S1), whereas SA pups did not differ from controls. Once pups reached the ventrum, time spent there was consistent across treatment groups (Fig. 1E), indicating that social adversity impaired the decision to initiate affiliative contact rather than the maintenance of contact itself.

This ventrum-specific deficit was accompanied by broader disruption of dam approach. SM pups were significantly less likely to approach the dam than BA controls (HR = 0.47, p = 0.003), whereas SA pups showed only a trend toward reduced approach (HR = 0.62, p = 0.055; Supp. Fig. 1A-B, Table S2), and SM pups spent significantly less time near the dam throughout the session, while SA pups again showed an intermediate phenotype (Supp. Fig. 1C). These differences were not explained by body weight, brain weight, brain-to-body mass ratio, or persistent locomotor differences (Supp. Fig. 2-4). Together, these findings indicate that pairing adversity with the maternal context produces substantially greater disruption of infant social approach than stress alone, with a specific impact on approach to the affiliative ventrum region.

To determine whether these early behavioral alterations persisted beyond infancy, DAM-exposed animals were tested in the two-choice social preference (2-CSP) task during adolescence (P45). Neither SM nor SA animals differed from controls in overall social or asocial zone occupancy (Fig. 1G, I). Instead, SM animals spent significantly more time in the neutral zone during the first 10 minutes of testing, with a trend toward greater neutral zone occupancy across the full session (Fig. 1H). Together, these findings indicate that adverse social experience produces persistent changes in social behavior that evolve from overt disruption of maternal approach in infancy to subtler alterations in social engagement during adolescence.

Sex-stratified analyses revealed that the behavioral consequences of social adversity differed between females and males. In infancy, both female and male SM pups approached the dam later and spent less time near her than their respective BA controls (Supp. Table 3,4; Supp. Fig. 5). However, disruption of affiliative ventrum approach was largely driven by females: SM-exposed females, but not males, approached the ventrum significantly later and spent less time near it than BA females, although direct sex comparisons within condition were not significant (Supp. Table 5,6; Supp. Fig. 5). In adolescence, this divergence became more pronounced. SM-exposed females spent significantly less time in the social zone and more time in the neutral zone than SA females, with a similar trend relative to BA females, whereas no significant differences were detected among males (Supp. Fig. 6). Thus, although adverse social experience altered social behavior in both sexes, females showed preferential disruption of affiliative social behavior across development, whereas males exhibited broader avoidance of the maternal stimulus during infancy without persistent deficits in adolescent social preference.

### Social adversity remodels behavior-associated BLA transcriptional networks during development

To identify central molecular correlates of the behavioral effects of adverse social experience, we performed bulk RNA sequencing of the basolateral amygdala (BLA) collected immediately following social behavior testing in infancy (P13) and adolescence (P45). Pairwise differential expression analyses identified significant transcriptional changes across adversity conditions at both developmental stages (Fig. 2A, C–E; Fig. 3A, C–E). However, principal component analysis revealed little separation between groups (Fig. 2B; Fig. 3B), suggesting that adversity remodels distributed transcriptional programs rather than producing large shifts in global gene expression.

**Figure 2.**
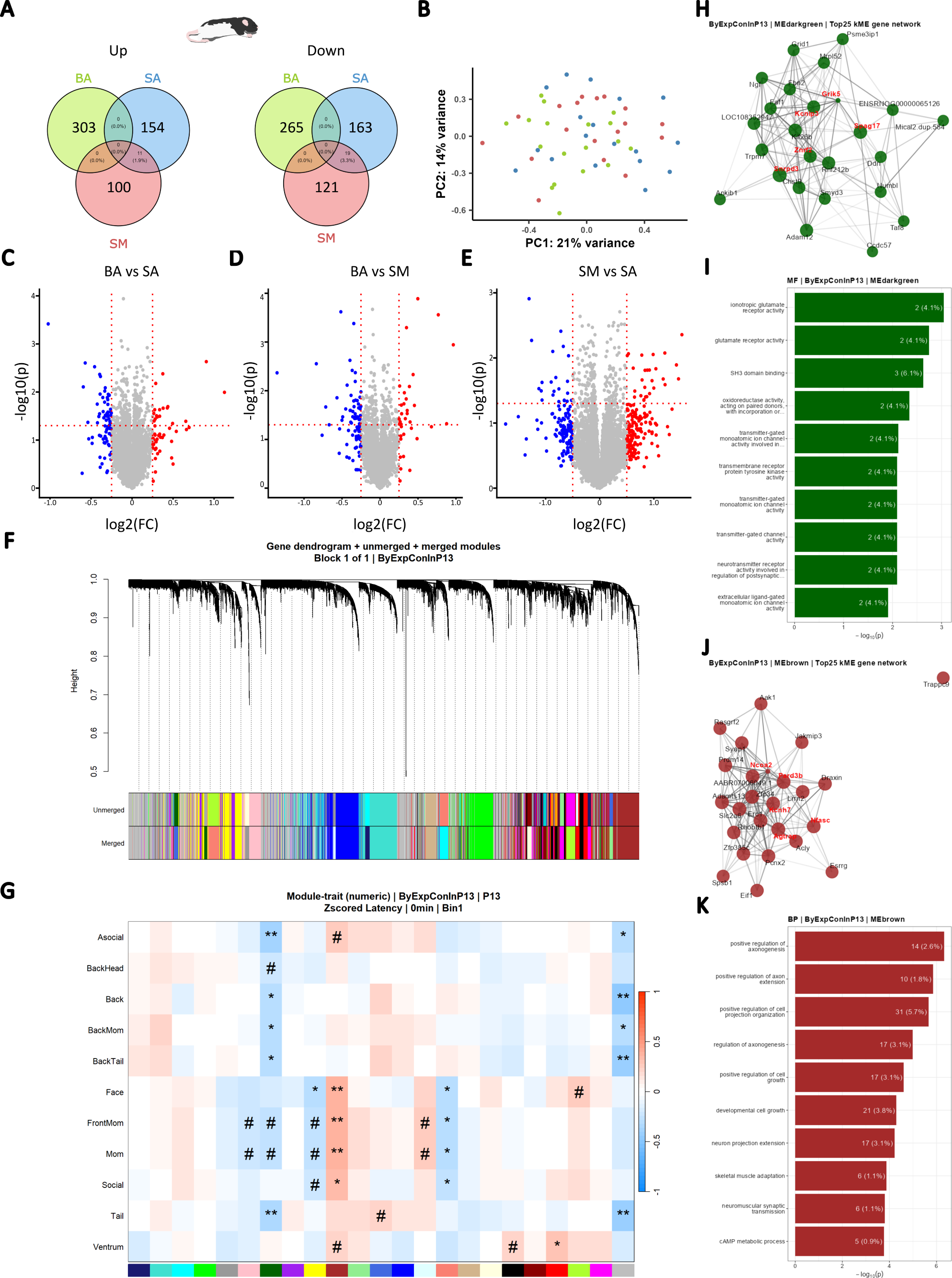
Effects of social and nonsocial ELS on the transcriptomic profile of the infant BLA. (A) Venn diagram of up (left) and down (right) regulated genes in infant BLA. Values represent number of genes up or down regulated in treatment conditions relative to BA. (B) Principal component analysis (PCA) of infant BA, SA, and SM BLA’s transcriptomic profile. (C, D, E) Volcano plots highlighting significantly up- and down-regulated genes for each pair-wise comparison. Left to right, BA vs SA, BA vs SM, and SM vs SA. −log10 of the pvalue is plotted on the y-axis, and log2 of the fold change (FC) between both conditions is plotted on the x-axis. Horizontal red dashed line drawn at Veritical red dashed lines drawn at −0.25 and −.25. Red and blue indicate a FC of ± 0.5, respectively. Gray indicates a FC between −0.5 and 0.5. (F) Weighted Gene CoExpression Network Analysis of infant (P13) samples reveals that genes in the BLA cluster into distinct modules. Each branch in the dendrogram represents a gene. Each gene is assigned to a cluster based on its distance to other genes (i.e., intramodular connectivity, kME), resulting in unmerged modules. Modules that are 75% correlated are merged. (G) Heatmap of correlations between merged modules and latency to approach infant pup data reveals novel module-trait relationships. Red and blue indicate positive and negative correlation, respectively. # indicates p < 0.1, * indicates p < 0.05, ** indicates p < 0.01, and *** indicates p < 0.001. (H, J) Network visualization of top 25 hub-genes in dark green (H) and brown (J) module by kME (a.k.a, module membership) graphed using Fruchterman-Reingold layout. Red text highlights the five hub-genes with the highest average kME. (I, K) Overrepresentation Analysis of genes in the dark green (I) and brown (K) modules reveal glutamate neurotransmission and axogenesis as key regulators of social behavior in infant BLA.

**Figure 3.**
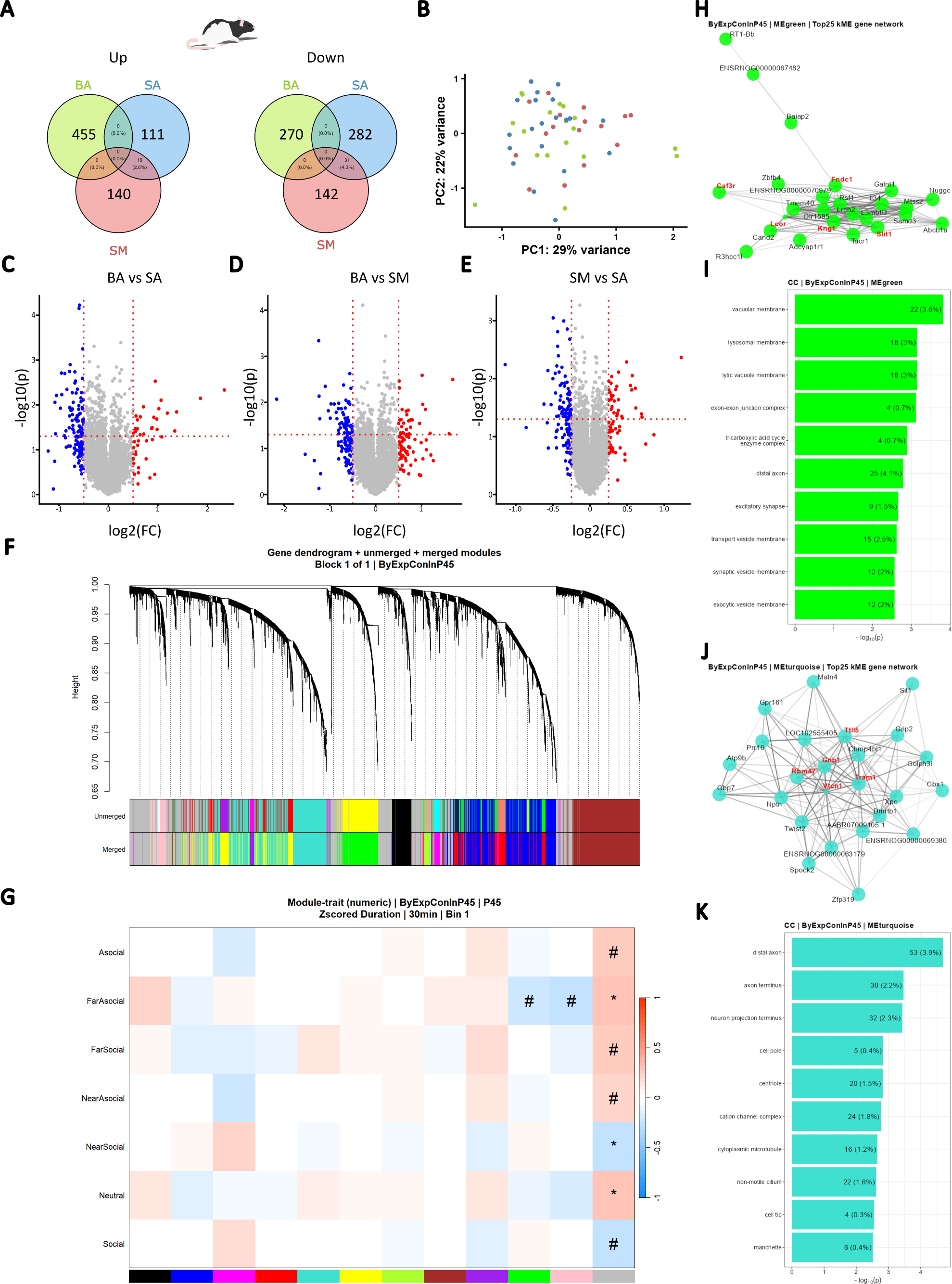
Effects of social and nonsocial ELS on the transcriptomic profile of the adolescent BLA. (A) Venn diagram of up (left) and down (right) regulated genes. Values represent number of genes up or down regulated in treatment conditions relative to BA. (B) PCA of adolescent BA, SA, and SM BLA’s transcriptomic profile. (C, D, E) Volcano plots highlighting significantly up- and down-regulated genes for each pair-wise comparison. Left to right, BA vs SA, BA vs SM, and SM vs SA. −log10 of the pvalue is plotted on the y-axis, and log2 of the fold change (FC) between both conditions is plotted on the x-axis. Horizontal red dashed line drawn at Veritical red dashed lines drawn at −0.25 and −.25. Red and blue indicate a FC of ± 0.5, respectively. Gray indicates a FC between −0.5 and 0.5. (F) Weighted Gene CoExpression Network Analysis of adolescent (P45) samples reveals that genes in the BLA cluster into distinct modules. Each branch in the dendrogram represents a gene. Each gene is assigned to a cluster based on its distance to other genes, resulting in unmerged modules. Modules that are 75% correlated are merged. (G) Heatmap of correlations between merged modules and time spent in (i.e., duration) pup data reveals novel module-trait relationships. Red and blue indicate positive and negative correlation, respectively. # indicates p < 0.1, * indicates p < 0.05, ** indicates p < 0.01, and *** indicates p < 0.001. (H, J) Network visualization of top 25 hub-genes in green (H) and turquoise (J) module by kME graphed using Fruchterman-Reingold layout. Red text highlights the five hub-genes with the highest average kME. (I, K) Overrepresentation Analysis of genes in the green (I) and turquoise (K) modules reveal that neither vacuolar membranes surrounding endosomes, lysosomes, and synaptic vesicles nor axons are associated with social behavior in adolescent BLA.

To identify coordinated transcriptional networks associated with behavior, we performed weighted gene co-expression network analysis (WGCNA). In the infant BLA, 26 gene modules were identified, several of which were significantly associated with maternal social approach behavior (Fig. 2F–G). The dark green module positively correlated with latency to approach the dam and maternal face, whereas the brown module showed a trend toward association with ventrum approach latency. Functional enrichment of highly connected hub genes implicated axonogenesis, axon extension, and projection organization within the dark green module and postsynaptic specialization and postsynaptic density within the brown module (Fig. 2H–K). These findings indicate that adverse social experience is associated with coordinated remodeling of developmental programs governing axonal growth and synaptic organization during infancy.

The transcriptional architecture shifted substantially by adolescence. WGCNA identified 11 gene modules in the adolescent BLA, with the green module negatively associated with both social zone occupancy and asocial-to-social transitions, whereas the turquoise module showed no relationship to social behavior (Fig. 3F–G). Hub genes within the green module were enriched for lysosomal and vacuolar membrane pathways, while the turquoise module was enriched for protein localization to the plasma membrane (Fig. 3H–K). Compared with infancy, these modules reflect a transition from transcriptional programs involved in circuit assembly toward pathways associated with membrane trafficking and cellular homeostasis, consistent with the changing developmental state of the adolescent BLA.

Because behavioral effects differed between females and males, we next asked whether adversity similarly produced sex-specific transcriptional organization (summarized in Supp. Table 7). In infancy, WGCNA identified 28 modules in females and 23 in males (Supp. Fig. 7,8). Female modules associated with maternal approach behavior were enriched for cholesterol transport and dendritic organization, whereas male modules implicated regulation of neurogenesis, nervous system development, glial differentiation, and translational control. Notably, cholesterol metabolism emerged only in female behavior-associated modules, linking central transcriptional organization to the bile acid pathway examined throughout this study.

Sex differences became even more pronounced in adolescence. Female WGCNA identified 20 modules whose expression strongly correlated with social and asocial zone occupancy and were enriched for synaptic and postsynaptic signaling pathways (Supp. Fig. 9,10). In contrast, no behavior-associated transcriptional modules out of 10 were detected in adolescent males. Together, these findings indicate that adverse social experience reorganizes coordinated BLA transcriptional networks across development and that the relationship between these networks and social behavior becomes increasingly sex-specific with maturation.

### Early social adversity alters developmental relationships between bile acid signaling and social behavior

To identify peripheral metabolic correlates of the behavioral effects of adverse social experience, we performed untargeted serum metabolomics immediately following social behavior testing in infancy (P13) and adolescence (P45). Pairwise analyses identified multiple metabolites that differed across adversity conditions at both developmental stages (Fig. 4A, C–E; Fig. 5A, C–E). In infancy, principal component analysis showed partial separation of SM animals from both BA and SA groups, whereas BA and SA animals largely overlapped (Fig. 4B), suggesting that adverse social experience produces a metabolomic profile distinct from stress exposure alone.

**Figure 4.**
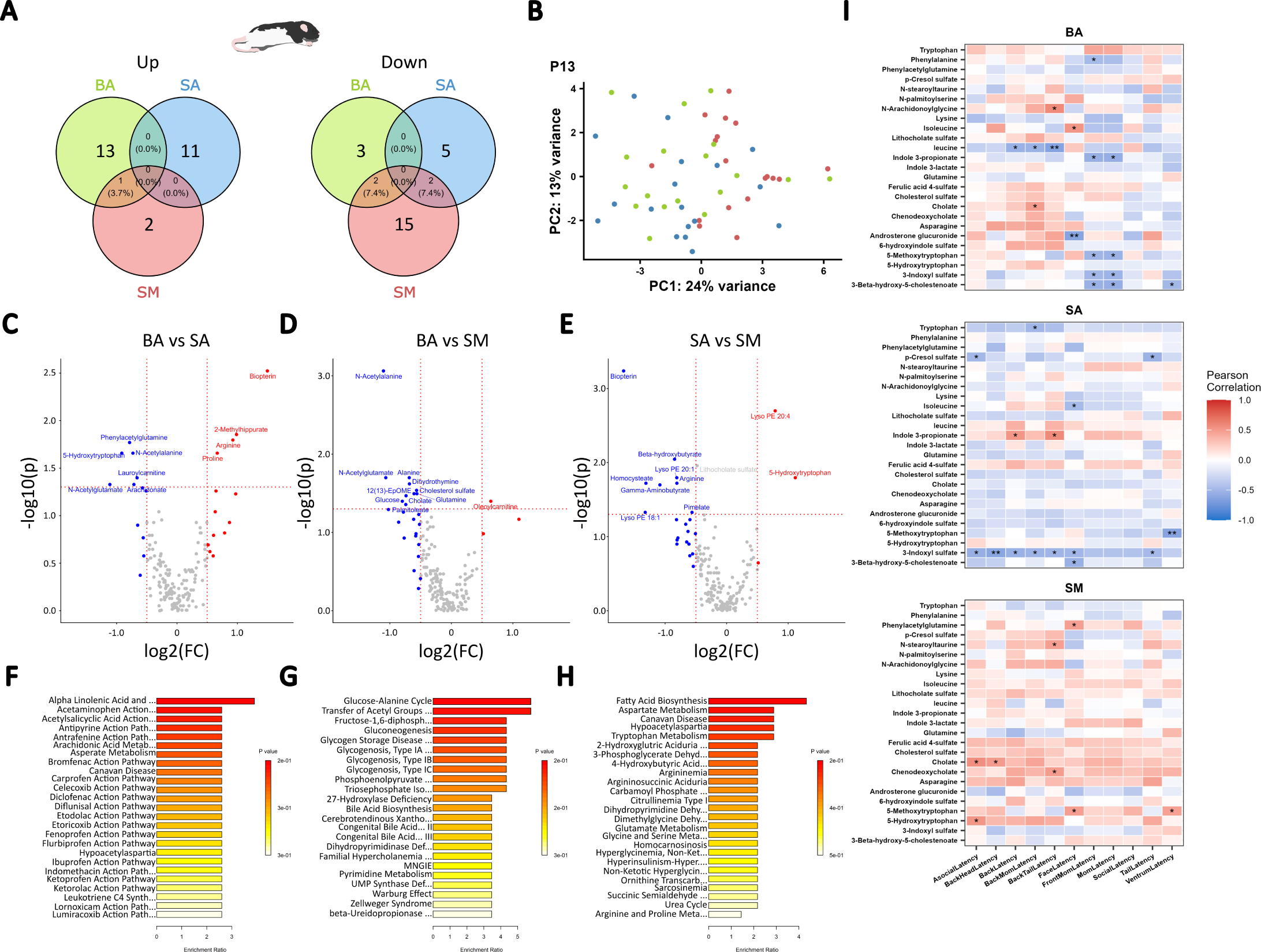
Social and non-social ELS differentially alter infant blood serum metabolome following MSA in infant pups. (A) Venn diagram of up (left) and down (right) regulated metabolites in infant blood serum. Values represent number of metabolites up or down regulated in each condition. (B) Principal component analysis of infant BA, SA, and SM’s blood serum metabolome. (C, D, E) Volcano plots highlighting significantly up- and down-regulated metabolites for each pair-wise comparison. Left to right, BA vs SA, BA vs SM, and SM vs SA. - log10 of the pvalue is plotted on the y-axis, and log2 of the fold change (FC) between both conditions is plotted on the x-axis. Horizontal red dashed line drawn at Veritical red dashed lines drawn at −0.25 and −.25. Red and blue indicate a FC of ± 0.5, respectively. Gray indicates a FC between −0.5 and 0.5. Labeled metabolites indicate a significant (p < 0.05) FC of ± 0.5. (F, G, H) Overrepresentation Analysis of all significantly different metabolites within BA vs SA (F), BA vs SM (G), and SA vs SM (H) does not reveal significantly overrepresented pathways. (I) Heatmap of metabolite-latency correlations reveal treatment dependent relationships. Red and blue indicate positive and negative correlation, respectively. * indicates p < 0.05, ** indicates p < 0.01, and *** indicates p < 0.001.

**Figure 5.**
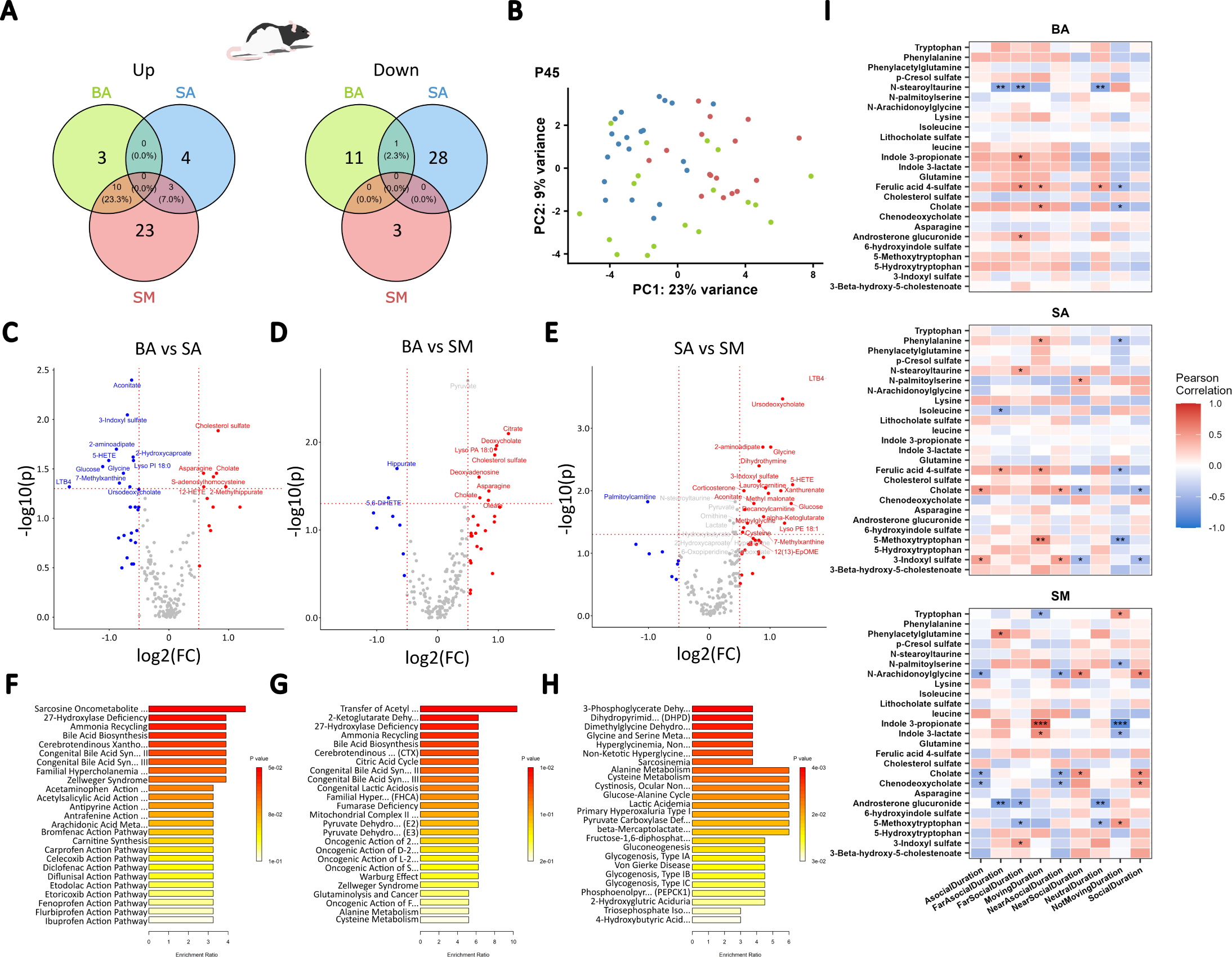
Social and non-social ELS alter bile and fatty acid plus tryptophan metabolism following 2-CSP in adolescent pups. (A) Venn diagram of up (left) and down (right) regulated metabolites in adolescent blood serum. Values represent number of metabolites up or down regulated in each condition. (B) PCA of adolescent BA, SA, and SM’s blood serum metabolome. (C, D, E) Volcano plots highlighting significantly up- and down-regulated metabolites for each pair-wise comparison. Left to right, BA vs SA, BA vs SM, and SM vs SA. −log10 of the pvalue is plotted on the y-axis, and log2 of the fold change (FC) between both conditions is plotted on the x-axis. Horizontal red dashed line drawn at Veritical red dashed lines drawn at −0.25 and −.25. Red and blue indicate a FC of ± 0.5, respectively. Gray indicates a FC between −0.5 and 0.5. Labeled metabolites indicate a significant (p < 0.05) FC of ± 0.5. (F, G, H) Overrepresentation Analysis of all significantly different metabolites within BA vs SA (F), BA vs SM (G), and SA vs SM (H) reveals that bile acid, fatty acid, and tryptophan metabolism are key targets of stress. (I) Heatmap of metabolite-duration correlations reveal treatment dependent relationships. Red and blue indicate positive and negative correlation, respectively. * indicates p < 0.05, ** indicates p < 0.01, and *** indicates p < 0.001.

Rather than converging on a single enriched pathway, infancy was characterized by altered relationships between circulating bile acid and tryptophan metabolites and social behavior. In control animals, several metabolites, including the CDCA precursor 3β-hydroxy-5-cholestenoate, indole-3-propionate, 5-methoxytryptophan, and 3-indoxyl sulfate were associated with faster maternal and ventrum approach (Fig. 4I). These relationships were altered following adversity. In SA animals, metabolite–behavior associations shifted toward non-social behavioral measures, whereas in SM animals several correlations involving bile acids and tryptophan metabolites were weakened or reversed. Notably, cholic acid became associated with approach to asocial regions, while 5-methoxytryptophan exhibited the opposite relationship with affiliative ventrum approach observed in controls. Although metabolite set enrichment analysis did not identify significantly enriched pathways in infancy (Fig. 4F–H), pairwise analyses implicated both bile acid and tryptophan metabolism, with alterations in cholic acid, cholesterol sulfate, and 5-hydroxytryptophan across adversity conditions (Fig. 4C–E). Together, these findings suggest that early social adversity alters the normal relationships between peripheral metabolism and infant social behavior before coordinated pathway-level changes emerge.

Sex-stratified metabolomic analyses revealed distinct developmental responses in females and males (Supp. Table 8; Supp. Fig. 11,12). Female infants exhibited broader alterations in bile acid and tryptophan metabolites than males, including reduced ursodeoxycholate following social adversity and reduced 5-hydroxytryptophan following non-social adversity. In contrast, males showed relatively limited metabolite changes, primarily involving lithocholate sulfate. Metabolite–behavior relationships also differed by sex. In control females, CDCA negatively correlated with maternal approach latency, whereas control males showed stronger associations with tryptophan metabolites. Following social adversity, females, but not males, developed positive correlations between cholic acid and delayed maternal and ventrum approach, paralleling the female-specific cholesterol-related transcriptional networks identified in the BLA.

By adolescence, metabolomic organization became more structured. Principal component analysis demonstrated clearer separation among adversity groups than in infancy (Fig. 5B), and metabolite set enrichment analysis identified significant alterations in bile acid biosynthesis, cholesterol metabolism, and amino acid pathways (Fig. 5F–H). Cholic acid and cholesterol sulfate remained altered following adversity, while corticosterone increased specifically following social adversity, indicating persistent endocrine and metabolic consequences of early adverse experience.

Behavior–metabolite relationships also differed from those observed in infancy. Cholic acid positively correlated with asocial behavior following SA but with social behavior following SM (Fig. 5I), indicating that early adversity changes the behavioral associations of bile acid signaling rather than simply altering metabolite abundance. Sex-stratified analyses again revealed distinct metabolic architectures (Supp. Table 8; Supp. Fig. 13,14). Female adolescents showed selective alterations in cholic acid and deoxycholate, whereas males exhibited broader disruption of bile acid, tryptophan, amino acid, and glucose metabolism together with significant pathway-level changes that were absent in females.

Together, these findings demonstrate that a single period of adverse social experience produces enduring alterations in peripheral metabolism that are expressed differently across development. Rather than simply changing metabolite abundance, early social adversity alters developmental relationships between bile acid signaling and social behavior, nominating bile acid signaling as a candidate mechanism linking adverse caregiving to disrupted social behavior and as a therapeutically tractable pathway for intervention.

### Chenodeoxycholic acid supplementation during early-life adversity selectively rescues infant social deficits

To determine whether pharmacologic targeting of bile acid signaling could rescue the behavioral consequences of adverse caregiving, pups exposed to Scarcity Adversity via Limited Bedding (SAM-LB) or control (Ctrl) rearing from P8–P12 received daily oral vehicle (Veh), CA, or CDCA throughout the adversity period and were subsequently assessed during MSA (P13– 15) and the 2-CSP task (P45–50) (Fig. 6A).

**Figure 6.**
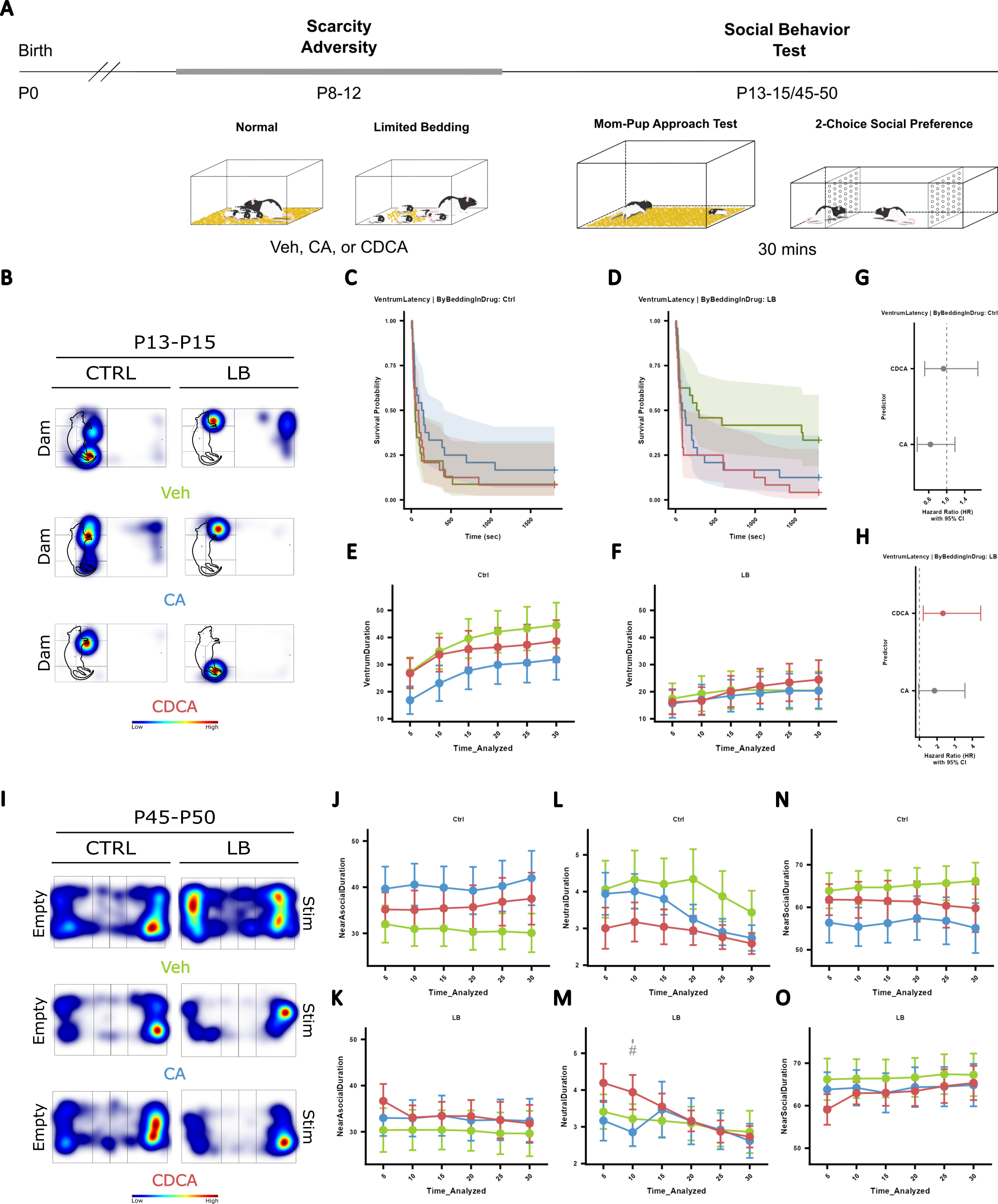
Primary bile acid, Cholic Acid (CA) and Chenodeoxycholic Acid (CDCA), supplementation during ELS prevents expression of social approach deficit. (A) Experimental design of primary bile acid supplementation during scarcity adversity via limited bedding (SAM-LB). Briefly, litters with six males and females are randomly assigned to control (CTRL) or limited bedding (LB) bedding conditions. Two males and females from each litter is then either is orally administered DMSO vehicle (Veh), 2.5 mg/kg CA, or 2.5 mg/kg CDCA. On P13-P15, infant social behavior is tested using the maternal social approach test, then on P45-P50 adolescent social behavior is tested using 2-CSP. (B) Representative heatmaps of Veh, CA, and CDCA treated Ctrl and LB infant pups in maternal social approach test show relative position of pups to an anesthetized maternal-odor producing dam (Dam). Dam outline overlaid onto heatmap. Blue and red indicate low and high proportion respectively. (C, D) Kaplan-Meier survival curves of Veh, CA, and CDCA treated Ctrl (C) and LB (D) exposed infant pups’ latency to approach dam. Survival probability (y-axis) of approaching the mom is graphed for each treatment as a function of Time (sec). (G, H) Cox-proportional hazard and 95% confidence interval (CI) of latency to approach the dam by CA and CDCA treated pups relative to Veh treated Ctrl (G) or LB (H) infant pups. Red indicates significantly different and gray indicates no significant difference. (E, F) Line graph of proportion of total duration (%) spent around the dam (i.e., mom) by the Veh, CA, and CDCA treated Ctrl (E) and LB (F) infant pups is plotted for each 5 min time bin. (I) Representative heatmaps of duration near empty or social stim (Stim) side of 2-CSP chamber by Veh, CA, and CDCA treated Ctrl and LB adolescent pups in 2-CSP. Blue and red indicate low and high proportion respectively. (J-O) Line graphs of proportion of total duration (%) spent near the social stim (NearSocial, left), neutral (middle), and near the empty (Empty, right) side of the 2-CSP apparatus are plotted within each 5 min time bin by the Veh, CA, and CDCA treated Ctrl (top) and LB (bottom) adolescent pups. Area under the curve of proportion of total duration spent around or within a region of interest is compared across each comparison (Veh vs CA, Veh vs CDCA, CA vs CDCA) and plotted to the right of the 30 min time bin value. Green is Veh, blue is CA, and red is CDCA. # indicates p < 0.1, * indicates p < 0.05, ** indicates p < 0.01, and *** indicates p < 0.001. Additionally, light gray * or # reflect comparison between BA and SA, black * or # reflect comparison between BA and SM, and dark gray * or # reflect comparison between SA and SM.

Because SAM-LB degrades the quality of maternal care without pairing the dam with an aversive stimulus, ventrum approach latency was designated as the primary behavioral outcome. In contrast to the Deconstructed Adversity Model (DAM), which produces conditioned avoidance of the dam as a whole, the behavioral consequences of SAM-LB were expected to be expressed most strongly in affiliative approach to the ventrum, where nursing and grooming occur. Consistent with this prediction and previous studies using this paradigm,^5,32^ SAM-LB produced no significant effect on overall dam approach latency (MomLatency), whereas vehicle-treated LB pups approached the ventrum significantly later than Ctrl Veh animals (*p* = 0.016), replicating our previous findings and confirming that SAM-LB selectively disrupts affiliative social approach rather than inducing generalized maternal avoidance (Fig. 6C–D; Supp. Table 9,10; Supp. Fig. 15). No treatment group differences were observed for growth milestones or locomotion (Supp. Fig. 16). The differential effects of DAM and SAM-LB on maternal behavior— broader disruption of dam approach in DAM versus selective impairment of ventrum approach in SAM-LB—are consistent with the distinct mechanisms by which the two models alter mother– infant interactions and further validate ventrum latency as the primary behavioral endpoint for the caregiving-quality manipulation.

CDCA, but not CA, rescued this deficit. Within the LB condition, CDCA-treated pups approached the ventrum significantly faster than LB Veh animals (HR lower bound = 1.22, *p* = 0.011; panel log-rank *p* = 0.031), whereas CA-treated LB pups showed only a non-significant trend toward improvement (*p* = 0.066) (Table 9; Fig. 6G–H). Importantly, neither CDCA nor CA altered ventrum approach in control-reared animals, indicating that rescue was specific to the adversity context rather than reflecting a general enhancement of social approach. Likewise, neither bedding condition nor drug treatment affected ventrum or dam approach duration, demonstrating that CDCA selectively influenced the decision to initiate affiliative contact rather than the quality or maintenance of social interaction once contact was established.

Sex-stratified analyses revealed that the SAM-LB-induced ventrum approach deficit was driven primarily by males. Vehicle-treated LB males approached the ventrum significantly later than Ctrl males, whereas LB females showed only a non-significant trend in the same direction (Supp. Table 12,13; Supp. Fig. 17). Although CDCA treatment reduced ventrum approach latency in both sexes, rescue did not reach statistical significance within either sex alone, and no sex-by-treatment interaction was detected (Supp. Table 13-18). Because the baseline behavioral deficit was substantially larger in males, the pooled rescue is most parsimoniously interpreted as reflecting rescue of the predominant male phenotype. Nevertheless, larger sex-balanced cohorts will be required to determine whether CDCA acts preferentially in males or produces comparable rescue in both sexes.

To determine whether these behavioral effects persisted beyond infancy, animals were evaluated during adolescent 2-CSP. Bedding condition alone did not produce a robust social preference deficit at this age (Supp. Fig. 18): no effects of bedding on time spent in the social, neutral, or asocial zones were detected, apart from a transient trend toward increased neutral zone occupancy in LB animals during the 10-minute time bin (Fig. 6I–O). Bile acid supplementation likewise had no effect on adolescent zone occupancy, and no sex-specific effects of bedding or treatment were observed (Supp. Fig. 19).

Together, these findings demonstrate that transient pharmacological manipulation of bile acid signaling during early-life adversity is sufficient to rescue the immediate behavioral consequences of adverse caregiving. The specificity of rescue for CDCA, but not the related primary bile acid CA, supports bile acid signaling as a therapeutically tractable pathway for modifying infant social behavior.

## DISCUSSION

The principal finding of this study is that transient pharmacologic manipulation of peripheral bile acid signaling during early-life adversity is sufficient to rescue an infant social behavioral deficit produced by adverse caregiving. Using complementary models of early-life adversity, we show that social, but not non-social adversity disrupts peripheral bile acid and tryptophan metabolism, reorganizes behavior-associated basolateral amygdala (BLA) transcriptional networks, and alters developmental relationships between bile acid signaling and social behavior. Most importantly, oral supplementation with chenodeoxycholic acid (CDCA), but not the related primary bile acid cholic acid (CA), selectively rescued the infant social behavioral deficit produced by caregiving scarcity. Together, these findings identify bile acid signaling as a mechanistically informative and pharmacologically tractable pathway linking adverse early social experience to disrupted social behavior. They further reveal a strikingly sex-dependent architecture of behavioral, metabolic, and transcriptomic responses to early adversity. In the Deconstructed Adversity Model (DAM), females exposed to social adversity exhibited preferential disruption of affiliative ventrum approach in infancy and reduced social engagement in adolescence, whereas males showed broader maternal avoidance. In contrast, in the Scarcity Adversity Model via Limited Bedding (SAM-LB), the infant ventrum approach deficit was predominantly observed in males and was selectively rescued by CDCA. Although the behavioral expression of vulnerability differed across models, peripheral metabolic and BLA transcriptional responses consistently showed sex-, age-, and adversity-type-dependent organization, suggesting that biological sex shapes how adverse social experience is translated into long-term changes in social behavior. To our knowledge, this study provides the first evidence that primary bile acid supplementation can rescue social behavioral deficits produced by adverse early caregiving and identifies bile acid signaling as a promising therapeutic target for modifying the developmental consequences of early-life adversity.

A major strength of the present study is the ability to distinguish the consequences of adverse social experience from those of stress alone. Using the Deconstructed Adversity Model, we found that social, but not non-social, adversity produced the most robust alterations in peripheral bile acid and tryptophan metabolism, paralleling the selective behavioral deficits observed across development. These findings extend previous work showing that early-life adversity alters the gut microbiome independently of maternal microbial transmission^33^ by demonstrating that adverse social experience also reshapes downstream bile acid metabolism in a sex-, age-, and adversity-type-dependent manner. The specificity of these metabolic alterations argues against a generalized stress response and instead supports the existence of a peripheral pathway selectively engaged by disrupted early social interactions. This distinction has important implications for understanding how different forms of adversity become biologically embedded and may help explain why adverse caregiving exerts particularly profound effects on later social behavior.

The selective rescue produced by CDCA provides the strongest evidence that bile acid signaling is not simply altered by early adversity but can be therapeutically targeted. Importantly, this effect was specific to CDCA; the structurally related primary bile acid cholic acid (CA) failed to produce significant behavioral improvement. Although the mechanism underlying this specificity remains to be determined, it is consistent with the greater potency of CDCA as an agonist of the farnesoid X receptor (FXR): CDCA is the most potent endogenous FXR agonist, whereas CA is among the weakest,^34–36^ providing a receptor-level basis for the compound specificity we observe and suggesting that receptor-mediated bile acid signaling contributes to the observed behavioral effects. These findings extend previous work in Chronic Unpredictable Mild Stress (CUMS) models, in which depression-like behavior is accompanied by altered bile acid metabolism and antidepressant treatment normalizes these abnormalities while increasing circulating CDCA.^22,37^ Likewise, direct administration of CDCA reduces depression-like behavior in rodents.^14^ Unlike these previous studies, however, our findings demonstrate that pharmacologic manipulation of bile acid signaling is sufficient to modify a developmental social behavioral consequence of adverse caregiving. Together, these convergent observations support bile acid signaling as a conserved pathway linking stress, peripheral metabolism, and behavioral dysfunction across distinct models of adversity while identifying CDCA as a promising candidate for intervention during early development.

Although the rescue experiment demonstrates that pharmacologic modulation of bile acid signaling is sufficient to modify infant social behavior, the transcriptomic and metabolomic analyses provide important clues regarding the biological pathways through which early adversity may exert these effects. Rather than producing widespread changes in global gene expression, social adversity preferentially reorganized coordinated BLA gene co-expression networks associated with social behavior. These network-level changes occurred in parallel with alterations in peripheral bile acid and tryptophan metabolism,^38^ suggesting coordinated remodeling of central and peripheral biological processes following adverse social experience. Notably, the strongest convergence was observed in females, in whom behavior-associated BLA transcriptional modules coincided with cholesterol- and bile acid-related metabolic signatures. Although these associations remain correlative, they support a model in which peripheral metabolic alterations accompany selective remodeling of neural circuits regulating social behavior rather than reflecting generalized consequences of stress. Consistent with the possibility that CDCA influences central nervous system function, recent work demonstrated that CDCA reduces infarct volume following ischemic stroke,^39^ suggesting that its biological actions extend beyond peripheral metabolism. Determining whether oral CDCA rescues social behavior through direct actions within the BLA, indirect modulation of peripheral metabolic signaling, or coordinated effects across both systems will be an important direction for future investigation.

The coordinated behavioral, metabolic, and transcriptomic responses observed in this study further suggest that biological sex shapes how adverse social experience is translated into long-term changes in social behavior. Importantly, our findings do not indicate that one sex is uniformly more vulnerable than the other. Rather, the behavioral consequences depended on both the nature of the adversity and the behavioral measure examined. In the DAM model, females showed greater disruption of affiliative ventrum approach during infancy and reduced social engagement during adolescence, whereas males exhibited broader maternal avoidance. In contrast, the caregiving-scarcity model (SAM-LB) produced a predominantly male ventrum approach deficit that was selectively rescued by CDCA. Although these cross-model differences require confirmation in larger, sex-balanced cohorts, they suggest that different forms of early adversity engage distinct biological processes whose behavioral expression is moderated by sex. Notably, the strongest evidence for sex-dependent organization emerged at the molecular level. In the DAM model, adolescent female BLA co-expression modules showed robust relationships with social behavior that were absent in males, while female-specific cholesterol and bile acid signatures paralleled these transcriptional changes. These findings suggest that sex differences may be expressed more consistently in the organization of peripheral and central biological responses than in behavior alone, where multiple compensatory processes may influence the observed phenotype.

Several, non-mutually exclusive mechanisms could account for the sex-dependent patterns observed in this study. First, different forms of early adversity may preferentially engage distinct neural circuits. Shock-conditioned social adversity in the DAM paradigm may recruit fear-learning networks, whereas caregiving scarcity in SAM-LB may preferentially disrupt affiliative motivational circuits, potentially explaining why females exhibited greater behavioral disruption in DAM while males showed greater vulnerability in SAM-LB. Second, sex hormones may modulate bile acid signaling during development. Estrogen suppresses hepatic FXR activity,^27,40^ raising the possibility that sex-dependent differences in FXR signaling alter the threshold at which peripheral bile acid dysregulation influences developing social circuits. Such interactions could contribute to the male-predominant behavioral deficit observed following SAM-LB while simultaneously permitting more extensive transcriptional remodeling in females. Third, developmental sex differences in gut microbiome composition and bile acid metabolism may generate distinct peripheral metabolic environments that differentially engage central signaling pathways.^41^ These mechanisms are not mutually exclusive and may interact throughout development. Because molecular measures were not collected in the SAM-LB cohort, the mechanisms underlying the behavioral rescue remain to be established. Future studies combining circuit-specific manipulation of FXR signaling, gonadal hormone manipulations, and direct comparisons of DAM and SAM-LB within the same experimental cohort will be required to determine how adversity type and biological sex interact to shape vulnerability to early-life adversity.

An additional consideration concerns the developmental scope of these effects. The caregiving-scarcity-induced deficit was evident in infancy but not in the adolescent 2-CSP assay, and CDCA rescued the infant deficit that was present. The absence of a detectable adolescent phenotype does not imply that the infant disruption lacks developmental significance. Early social interactions provide critical opportunities for caregiver engagement, social learning, and the acquisition of behavioral competencies, and even transient disruptions in affiliative behavior during infancy may influence developmental trajectories in ways not captured by later behavioral assays.^42^ Consistent with the age-dependent expression of limited-bedding phenotypes, threat-related behavioral abnormalities in this model can be transiently masked during juvenility before re-emerging in adolescence,^43^ underscoring that the presence or absence of a phenotype at any single age reflects both the developmental stage and the specific behavior assayed. Whether early modulation of bile acid signaling influences later-emerging outcomes—particularly in adversity models that do produce adolescent phenotypes— will require longitudinal assessment and is an important direction for future work.

The identification of bile acid signaling as a modifiable pathway linking adverse social context to disrupted social behavior expands current understanding of how early-life adversity becomes biologically embedded. Rather than viewing peripheral metabolic changes as downstream consequences of stress, our findings support the possibility that they actively participate in shaping developmental trajectories of social behavior through a bile acid-brain-behavior axis. Because CDCA is already FDA-approved for human use, these findings provide an initial rationale for investigating bile acid–based interventions during sensitive developmental periods. More broadly, this work highlights the importance of integrating peripheral metabolism with circuit-level analyses to understand how early experiences become embedded in the developing brain and influence psychiatric risk across the lifespan.

## Materials and Methods

### Animals and adversity models

Male and female Long Evans rats were bred in-house, with cross-fostering and litter-based assignment used to control for litter effects. Two complementary early-life adversity paradigms were applied from postnatal day (P) 8 to P12. In the Deconstructed Adversity Model (DAM), pups were assigned to beaker-alone control (BA), repeated shock alone (non-social adversity, SA), or the same shock delivered in the presence of an anesthetized maternal-odor-producing dam (social adversity, SM). In the Scarcity Adversity Model via Limited Bedding (SAM-LB), bedding was reduced to 100 cc (LB) or maintained at 2000 cc (Ctrl), and maternal and pup behaviors were scored from daily recordings using BORIS.^44^

### Bile acid supplementation

SAM-LB and control pups received daily oral vehicle, cholic acid (CA; 2.5 mg/kg), or chenodeoxycholic acid (CDCA; 2.5 mg/kg) from P8 to P12.

### Behavioral testing

Infant pups (P13–15) underwent a 30-minute maternal social approach (MSA) test with an unfamiliar anesthetized maternal-odor-producing dam, during which latency to, time near, and visits to defined body-region zones—including the affiliative ventrum where nursing and grooming occur—were tracked. Adolescent pups (P45–50) underwent a 30-minute two-choice social preference (2-CSP) test in a three-chamber apparatus, with time near the social, neutral, and asocial zones tracked in EthoVision.^45^

### Tissue collection and molecular profiling

Serum and whole brain were collected 60 minutes after behavioral testing. The basolateral amygdala (BLA) was harvested by cryostat punch using published stereotaxic coordinates.^46^ Untargeted serum metabolomics was performed by ultrafast liquid chromatography–high-resolution tandem mass spectrometry (TripleTOF 5600), yielding 173 reliably quantified metabolites. BLA transcriptomes were profiled by bulk mRNA sequencing (Illumina NovaSeq); reads were processed with a Trimgalore–HISAT2– featureCounts pipeline^47–49^ and differential expression and network analyses used DESeq2^50^ and weighted gene co-expression network analysis (WGCNA).^51^

### Statistical analysis

Group comparisons used non-parametric tests (Wilcoxon rank-sum and Welch’s ANOVA). Censored approach-latency data were analyzed by Kaplan–Meier estimation with log-rank tests and Cox proportional-hazards models. Analyses were performed on pooled data, within each sex, and across sexes, and metabolomic and transcriptomic comparisons in the DAM cohort were not corrected for multiple comparisons given the adversity×sex×age design; significance was defined as p < 0.05. Full experimental and analytical details are provided in Supplementary Methods.

### Resource Availability

Further information and requests for resources and reagents should be directed to the lead contact, Maya Opendak.

#### Materials availability

This study did not generate new unique reagents.

#### Data and code availability

Raw bulk mRNA sequencing data have been deposited at NCBI GEO and are publicly available as of the date of publication (accession number [to be assigned]). Raw metabolomics data have been deposited at Metabolomics Workbench and are publicly available as of the date of publication (accession number [to be assigned]). All original code used for behavioral data aggregation, statistical analyses, and figure generation has been deposited at GitHub and is publicly available as of the date of publication (repository URL [to be assigned]). Any additional information required to reanalyze the data reported in this paper is available from the lead contact upon request.

## Acknowledgments

This work was supported by the National Institutes of Health through grants R00MH124434 and R01MH133456 to MO, R01MH131469 and R01MH131219 to NH, and by core support from the Intellectual and Developmental Disabilities Research Center (IDDRC) at Kennedy Krieger Institute, Johns Hopkins University (P50HD105328; U54HD079123) and the Integrated Genomics Center. Additional support was provided by the Kennedy Krieger Institute Goldstein Award, and the Johns Hopkins University Discovery Award (MO). Figures were created using custom R code.

## Author Contributions

C.M.: Conceptualization, Methodology, Investigation, Formal analysis, Writing – original draft. P.D.: Investigation (metabolomics), N.H.: Investigation (metabolomics), Funding acquisition. V.W., N.I., I.M., M.R., E.A.: Investigation. M.O.: Conceptualization, Supervision, Project administration, Funding acquisition, Writing – review & editing.

