## Supplementary Tables for "Bile acid signaling as a therapeutically tractable pathway linking early caregiving adversity to social behavior"

**Contents**

**Supplementary Table 1.** Log-rank test comparing hazard ratios (HR) of SA and SM treated infant pups’ latency to approach the ventrum against their BA counterparts.

**Supplementary Table 2.** Log-rank test comparing hazard ratios (HR) of SA and SM treated infant pups’ latency to approach the mom (any region) against their BA counterparts.

**Supplementary Table 3.** Log-rank test comparing hazard ratios (HR) of female and male SA and SM treated infant pups’ latency to approach the Dam against their BA counterparts.

**Supplementary Table 4.** Log-rank test comparing hazard ratios (HR) of BA, SA and SM treated infant Female pups’ latency to approach the Dam against their Male counterparts.

**Supplementary Table 5.** Log-rank test comparing hazard ratios (HR) of female and male SA and SM treated infant pups’ latency to approach the ventrum against their BA counterparts.

**Supplementary Table 6.** Log-rank test comparing hazard ratios (HR) of BA, SA and SM treated infant Female pups’ latency to approach the ventrum against their Male counterparts.

**Supplementary Table 7.** Sex- and condition-dependent BLA transcriptomic (WGCNA/GSEA) changes and module–behavior correlations across development

**Supplementary Table 8.** Sex- and condition-specific metabolomic changes and metabolite–behavior correlations across development

**Supplementary Table 9.** Log-rank test comparing hazard ratios (HR) of LB exposed infant pups’ latency to approach the ventrum against their Ctrl counterparts within each treatment condition.

**Supplementary Table 10.** Log-rank test comparing hazard ratios (HR) of CA and CDCA treated infant pups’ latency to approach the ventrum against their Veh counterparts within each bedding condition.

**Supplementary Table 11.** Log-rank test comparing hazard ratios (HR) of LB exposed infant pups’ latency to approach the Dam against their Ctrl counterparts within each drug treatment.

**Supplementary Table 12.** Log-rank test comparing hazard ratios (HR) of CA and CDCA treated infant pups’ latency to approach the Dam against their Veh counterparts within each bedding condition.

**Supplementary Table 13.** Log-rank test comparing hazard ratios (HR) of Female and Male Ctrl and LB exposed infant pups’ latency to approach the ventrum against their Veh treated counterparts.

**Supplementary Table 14.** Log-rank test comparing hazard ratios (HR) of Female and Male Veh, CA, and CDCA treated infant pups’ latency to approach the ventrum against their Ctrl counterparts.

**Supplementary Table 15.** Log-rank test comparing hazard ratios (HR) of Veh, CA, and CDCA treated Ctrl and LB infant Female (F) pups’ latency to approach the ventrum against their Male (M) counterparts.

**Supplementary Table 16.** Log-rank test comparing hazard ratios (HR) of Female and Male Ctrl and LB exposed infant pups’ latency to approach the Dam against their Veh treated counterparts.

**Supplementary Table 17.** Log-rank test comparing hazard ratios (HR) of Female and Male Veh, CA, and CDCA treated infant pups’ latency to approach the ventrum against their Ctrl counterparts.

**Supplementary Table 18.** Log-rank test comparing hazard ratios (HR) of Veh, CA, and CDCA treated Ctrl and LB infant Female (F) pups’ latency to approach the Dam against their Male (M) counterparts.

| Characteristic^1^ | Median (95% CI) | HR | 95% CI | p-value |
| --- | --- | --- | --- | --- |
| Condition |  |  |  |  |
| BA | 253 (178, 501) | — | — |  |
| SA | 187 (128, 320) | 1.07 | 0.64, 1.78 | 0.8 |
| SM | 1,043 (505, 1,694) | 0.53 | 0.31, 0.91 | 0.022 |
| ^1^Global Log-Rank Test p-value: p=0.020 | | | | |
| Abbreviations: CI = Confidence Interval, HR = Hazard Ratio | | | | |

Supplementary Table 1. Log-rank test comparing hazard ratios (HR) of SA and SM treated infant pups’ latency to approach the ventrum against their BA counterparts. Median latency along with the 95% confidence interval (CI) was calculated for all pups. HR, 95% CI of HR, and p-value was calculated for each comparison (BA vs SA and BA vs SM).

|  | Descriptive Summary | Cox Model | | |
| --- | --- | --- | --- | --- |
| Characteristic^1^ | Median (95% CI) | HR | 95% CI | p-value |
| Condition |  |  |  |  |
| BA | 28 (18, 97) | — | — |  |
| SA | 44 (24, 118) | 0.62 | 0.38, 1.01 | 0.055 |
| SM | 107 (34, 277) | 0.47 | 0.29, 0.78 | 0.003 |
| ^1^Global Log-Rank Test p-value: p=0.010 | | | | |
| Abbreviations: CI = Confidence Interval, HR = Hazard Ratio | | | | |

Supplementary Table 2. Log-rank test comparing hazard ratios (HR) of SA and SM treated infant pups’ latency to approach the mom (any region) against their BA counterparts. Median latency along with the 95% confidence interval (CI) was calculated for all pups. HR, 95% CI of HR, and p-value was calculated for each comparison (BA vs SA and BA vs SM).

|  | Descriptive Summary | Cox Model | | |
| --- | --- | --- | --- | --- |
| Characteristic | Median (95% CI) | HR | 95% CI | p-value |
| Female^1^ |  |  |  |  |
| BA | 46 (15, 264) | — | — |  |
| SA | 50 (36, 588) | 0.66 | 0.33, 1.31 | 0.2 |
| SM | 185 (53, 982) | 0.50 | 0.25, 1.00 | 0.050 |
| Male^2^ |  |  |  |  |
| BA | 22 (18, 100) | — | — |  |
| SA | 24 (23, 267) | 0.60 | 0.29, 1.24 | 0.2 |
| SM | 87 (29, 479) | 0.45 | 0.22, 0.93 | 0.032 |
| ^1^Log-Rank Test p-value (Within Female): p=0.13  ^2^Log-Rank Test p-value (Within Male): p=0.086 | | | | |
| Abbreviations: CI = Confidence Interval, HR = Hazard Ratio | | | | |

Supplementary Table 3. Log-rank test comparing hazard ratios (HR) of female and male SA and SM treated infant pups’ latency to approach the Dam against their BA counterparts. Median latency along with the 95% confidence interval (CI) was calculated for all pups. HR, 95% CI of HR, and p-value was calculated for each comparison (BA vs SA and BA vs SM).

|  | Descriptive Summary | Cox Model | | |
| --- | --- | --- | --- | --- |
| Characteristic | Median (95% CI) | HR | 95% CI | p-value |
| BA^1^ |  |  |  |  |
| Male | 22 (18, 100) | — | — |  |
| Female | 46 (15, 264) | 0.62 | 0.30, 1.28 | 0.2 |
| SA^2^ |  |  |  |  |
| Male | 24 (23, 267) | — | — |  |
| Female | 50 (36, 588) | 0.77 | 0.38, 1.57 | 0.5 |
| SM^3^ |  |  |  |  |
| Male | 87 (29, 479) | — | — |  |
| Female | 185 (53, 982) | 0.83 | 0.41, 1.68 | 0.6 |
| ^1^Log-Rank Test p-value (Within BA): p=0.2  ^2^Log-Rank Test p-value (Within SA): p=0.5  ^3^Log-Rank Test p-value (Within SM): p=0.6 | | | | |
| Abbreviations: CI = Confidence Interval, HR = Hazard Ratio | | | | |

Supplementary Table 4. Log-rank test comparing hazard ratios (HR) of BA, SA and SM treated infant Female pups’ latency to approach the Dam against their Male counterparts. Median latency along with the 95% confidence interval (CI) was calculated for all pups. HR, 95% CI of HR, and p-value was calculated for each comparison (BA vs SA and BA vs SM).

|  | Descriptive Summary | Cox Model | | |
| --- | --- | --- | --- | --- |
| Characteristic | Median (95% CI) | HR | 95% CI | p-value |
| Female^1^ |  |  |  |  |
| BA | 270 (98, 1,580) | — | — |  |
| SA | 292 (183, 1,548) | 0.99 | 0.49, 2.04 | >0.9 |
| SM | 1,420 (980, —) | 0.42 | 0.19, 0.91 | 0.029 |
| Male^2^ |  |  |  |  |
| BA | 235 (178, 681) | — | — |  |
| SA | 145 (72, 411) | 1.24 | 0.60, 2.57 | 0.6 |
| SM | 631 (201, —) | 0.69 | 0.33, 1.45 | 0.3 |
| ^1^Log-Rank Test p-value (Within Female): p=0.041  ^2^Log-Rank Test p-value (Within Male): p=0.3 | | | | |
| Abbreviations: CI = Confidence Interval, HR = Hazard Ratio | | | | |

Supplementary Table 5. Log-rank test comparing hazard ratios (HR) of female and male SA and SM treated infant pups’ latency to approach the ventrum against their BA counterparts. Median latency along with the 95% confidence interval (CI) was calculated for all pups. HR, 95% CI of HR, and p-value was calculated for each comparison (BA vs SA and BA vs SM).

|  | Descriptive Summary | Cox Model | | |
| --- | --- | --- | --- | --- |
| Characteristic | Median (95% CI) | HR | 95% CI | p-value |
| BA^1^ |  |  |  |  |
| Male | 235 (178, 681) | — | — |  |
| Female | 270 (98, 1,580) | 0.98 | 0.48, 2.01 | >0.9 |
| SA^2^ |  |  |  |  |
| Male | 145 (72, 411) | — | — |  |
| Female | 292 (183, 1,548) | 0.79 | 0.38, 1.63 | 0.5 |
| SM^3^ |  |  |  |  |
| Male | 631 (201, —) | — | — |  |
| Female | 1,420 (980, —) | 0.55 | 0.25, 1.21 | 0.14 |
| ^1^Log-Rank Test p-value (Within BA): p>0.9  ^2^Log-Rank Test p-value (Within SA): p=0.5  ^3^Log-Rank Test p-value (Within SM): p=0.13 | | | | |
| Abbreviations: CI = Confidence Interval, HR = Hazard Ratio | | | | |

Supplementary Table 6. Log-rank test comparing hazard ratios (HR) of BA, SA and SM treated infant Female pups’ latency to approach the ventrum against their Male counterparts. Median latency along with the 95% confidence interval (CI) was calculated for all pups. HR, 95% CI of HR, and p-value was calculated for each comparison (BA vs SA and BA vs SM).

| AGE | SEX | CONDITION | SIGNIFICANT WGCNA MODULE / GSEA PATHWAY RESULTS | MODULE–BEHAVIOR CORRELATION |
| --- | --- | --- | --- | --- |
| INFANT (P13) — POOLED | | | | |
| P13 | Pooled | BA vs SA vs SM | Significant genes up- and down-regulated across all conditions; PCA showed minimal clustering by condition | — |
| P13 | Pooled | All conditions (WGCNA) | 26 modules identified. Yellow module: positive regulation of projection organization, axogenesis, axon extension, and cell growth (GSEA). Turquoise module: postsynaptic specialization, neuron-to-neuron synapse, postsynaptic density, and proteasome regulatory particles (GSEA) | Yellow (+) ~ mom & face latency   Turquoise (trend) ~ ventrum latency |
| INFANT (P13) — SEX-SPECIFIC | | | | |
| P13 | Female | All conditions (WGCNA) | 28 modules identified. Royal blue module: stereocilia and dendritic cytoplasm (GSEA). Yellow module: regulation of sterol, cholesterol, and cation-transmembrane transport (GSEA). | Royal blue (+) ~ mom latency, incl. asocial (back, back of head, back of tail) latency   Yellow (−) ~ asocial region latency |
| P13 | Male | All conditions (WGCNA) | 23 modules identified. Light-yellow module: negative regulation of neurogenesis, nervous system development, and glial cell differentiation (GSEA). Dark turquoise module: translation initiation via RNA–transcription factor binding and DNA regulation (GSEA). | Light-yellow (+) ~ mom latency, incl. ventrum   Dark turquoise (−) ~ ventrum latency (amongst others) |
| ADOLESCENT (P45) — POOLED | | | | |
| P45 | Pooled | BA vs SA vs SM | Significant genes up- and down-regulated across all conditions; PCA showed no clustering by condition | — |
| P45 | Pooled | All conditions (WGCNA) | 11 modules identified. Green module: vacuolar membranes, lysosomal membranes, and lytic vacuole membranes (GSEA). Turquoise module: roof of mouth development, protein localization to cell periphery, and protein localization to plasma membrane (GSEA) | Green (−) ~ asocial-social transitions; far social & social duration.   Turquoise: not associated with social behavior |
| ADOLESCENT (P45) — SEX-SPECIFIC | | | | |
| P45 | Female | All conditions (WGCNA) | 20 modules identified (module-trait correlation). Pink module: synapse (GSEA); Yellow module: post-synapse (GSEA) | Green-yellow (−) ~ social zone time, (+) ~ asocial zone time; Tan (+) ~ social zone time, (−) ~ asocial zone time |
| P45 | Male | All conditions (WGCNA) | 10 modules identified; no significantly enriched GSEA pathways reported | No modules significantly correlated with time-spent behaviors |

Supplemental Table 7. Sex- and condition-dependent BLA transcriptomic (WGCNA/GSEA) changes and module–behavior correlations across development

*WGCNA module eigengenes (ME) were correlated with social behavior latency (infant MSA) or time-spent measures (adolescent 2-CSP). Only modules with significant (or trending) module-trait correlations are shown, with associated GSEA-enriched pathways/cellular compartments. “—” denotes not significant, not applicable, or not reported. Derived from Figs 2–3 and sex-stratified supplemental analyses.*

| AGE | SEX | CONDITION | SIGNIFICANT METABOLITE / PATHWAY CHANGE (vs BA unless noted) | METABOLITE–BEHAVIOR CORRELATION |
| --- | --- | --- | --- | --- |
| INFANT (P13) — POOLED | | | | |
| P13 | Pooled | SA vs SM | CA ↑ and cholesterol sulfate ↑ in SA vs SM | — |
| P13 | Pooled | SA | 5-Hydroxytryptophan ↑ | 3-Beta-hydroxy-5-cholestenoate ~ face latency; Trp metabolites shift to asocial zones |
| P13 | Pooled | SM | 5-Hydroxytryptophan ↓ | CA (−) ~ asocial latency; 5-Methoxytryptophan (+) ~ ventrum & face latency |
| P13 | Pooled | BA (control) | — | 3-Beta-hydroxy-5-cholestenoate (−) ~ mom & ventrum latency; Trp metabolites (−) ~ latency |
| INFANT (P13) — SEX-SPECIFIC | | | | |
| P13 | Female | SA | 5-Hydroxytryptophan ↓ | — |
| P13 | Female | SM | Ursodeoxycholate ↓ | CA (+) ~ mom & ventrum latency |
| P13 | Female | BA (control) | — | CDCA (−) ~ social approach latency |
| P13 | Male | SA vs SM | Lithocholate sulfate ↑ in SA vs SM | — |
| P13 | Male | SM | (no other significant BA/Trp changes) | CDCA (+) ~ mom latency |
| P13 | Male | BA (control) | — | Trp metabolites ~ latency (BA/cholesterol metabolites n.s.) |
| ADOLESCENT (P45) — POOLED | | | | |
| P45 | Pooled | SA & SM | CA ↑ and cholesterol sulfate ↑ (both vs BA) | CA (+) ~ asocial zone time (SA); CA (+) ~ social zone time (SM) |
| P45 | Pooled | SM vs SA | Corticosterone ↑ in SM vs SA | — |
| P45 | Pooled | Pathways | BA biosynthesis & cholesterol metab. enriched (BA vs SA, BA vs SM); amino-acid metab. enriched (SA vs SM) | — |
| ADOLESCENT (P45) — SEX-SPECIFIC | | | | |
| P45 | Female | SA | Cholate ↑ | BA & Trp metabolites ~ movement & neutral zone (condition-specific) |
| P45 | Female | SM | Deoxycholate ↑ | (as above) |
| P45 | Female | Pathways | No significant enrichment | — |
| P45 | Male | SA | 3-Indoxyl sulfate ↓, Ursodeoxycholate ↓; amino-acid & glucose metab. pathways ↓ (vs BA and SM) | Correlations sparse |
| P45 | Male | SM | Cholate ↑, Ursodeoxycholate ↑, 3-Indoxyl sulfate ↑, Pyruvate ↑, Corticosterone ↑ | Indole-3-propionate (+) ~ movement |

Supplemental Table 8. Sex- and condition-specific metabolomic changes and metabolite–behavior correlations across development

*Significant metabolite changes are relative to BA (control) unless a specific contrast is noted. Arrows indicate direction of change (↑ elevated, ↓ reduced). Correlations show the sign (+/−) of the Pearson relationship between metabolite level and the indicated behavioral latency or zone measure. “~” denotes “correlated with.” n.s. = not significant. Derived from Fig 4, Fig 5, and sex-stratified supplemental analyses.*

|  | Descriptive Summary | Cox Model | | |
| --- | --- | --- | --- | --- |
| Characteristic | Median (95% CI) | HR | 95% CI | p-value |
| Veh^1^ |  |  |  |  |
| Ctrl | 50 (37, 131) | — | — |  |
| LB | 280 (57, —) | 0.44 | 0.23, 0.86 | 0.016 |
| CA^2^ |  |  |  |  |
| Ctrl | 143 (46, 418) | — | — |  |
| LB | 103 (50, 272) | 1.17 | 0.63, 2.15 | 0.6 |
| CDCA^3^ |  |  |  |  |
| Ctrl | 77 (33, 154) | — | — |  |
| LB | 75 (43, 99) | 1.03 | 0.57, 1.87 | >0.9 |
| ^1^Log-Rank Test p-value (Within Veh): p=0.014  ^2^Log-Rank Test p-value (Within CA): p=0.6  ^3^Log-Rank Test p-value (Within CDCA): p>0.9 | | | | |
| Abbreviations: CI = Confidence Interval, HR = Hazard Ratio | | | | |

Supplementary Table 9. Log-rank test comparing hazard ratios (HR) of LB exposed infant pups’ latency to approach the ventrum against their Ctrl counterparts within each treatment condition. Median latency along with the 95% confidence interval (CI) was calculated for all pups. HR, 95% CI of HR, and p-value was calculated for Ctrl vs LB.

|  | Descriptive Summary | Cox Model | | |
| --- | --- | --- | --- | --- |
| Characteristic | Median (95% CI) | HR | 95% CI | p-value |
| Ctrl^1^ |  |  |  |  |
| Veh | 50 (37, 131) | — | — |  |
| CA | 143 (46, 418) | 0.64 | 0.34, 1.18 | 0.2 |
| CDCA | 77 (33, 154) | 0.93 | 0.51, 1.69 | 0.8 |
| LB^2^ |  |  |  |  |
| Veh | 280 (57, —) | — | — |  |
| CA | 103 (50, 272) | 1.85 | 0.96, 3.58 | 0.066 |
| CDCA | 75 (43, 99) | 2.33 | 1.22, 4.47 | 0.011 |
| ^1^Log-Rank Test p-value (Within Ctrl): p=0.3  ^2^Log-Rank Test p-value (Within LB): p=0.031 | | | | |
| Abbreviations: CI = Confidence Interval, HR = Hazard Ratio | | | | |

Supplementary Table 10. Log-rank test comparing hazard ratios (HR) of CA and CDCA treated infant pups’ latency to approach the ventrum against their Veh counterparts within each bedding condition. Median latency along with the 95% confidence interval (CI) was calculated for all pups. HR, 95% CI of HR, and p-value was calculated for Veh vs CA and Veh vs CDCA.

|  | Descriptive Summary | Cox Model | | |
| --- | --- | --- | --- | --- |
| Characteristic | Median (95% CI) | HR | 95% CI | p-value |
| Veh^1^ |  |  |  |  |
| Ctrl | 21 (12, 35) | — | — |  |
| LB | 16 (8.1, 34) | 1.00 | 0.56, 1.81 | >0.9 |
| CA^2^ |  |  |  |  |
| Ctrl | 18 (9.8, 39) | — | — |  |
| LB | 9.0 (5.4, 25) | 1.61 | 0.90, 2.87 | 0.11 |
| CDCA^3^ |  |  |  |  |
| Ctrl | 25 (13, 55) | — | — |  |
| LB | 7.5 (5.0, 34) | 1.58 | 0.88, 2.86 | 0.13 |
| ^1^Log-Rank Test p-value (Within Veh): p>0.9  ^2^Log-Rank Test p-value (Within CA): p=0.10  ^3^Log-Rank Test p-value (Within CDCA): p=0.12 | | | | |
| Abbreviations: CI = Confidence Interval, HR = Hazard Ratio | | | | |

Supplementary Table 11. Log-rank test comparing hazard ratios (HR) of LB exposed infant pups’ latency to approach the Dam against their Ctrl counterparts within each drug treatment. Median latency along with the 95% confidence interval (CI) was calculated for all pups. HR, 95% CI of HR, and p-value was calculated for Ctrl vs LB.

|  | Descriptive Summary | Cox Model |
| --- | --- | --- |

|  | Descriptive Summary | Cox Model | | |
| --- | --- | --- | --- | --- |
| Characteristic | Median (95% CI) | HR | 95% CI | p-value |
| Ctrl^2^ |  |  |  |  |
| Veh | 21 (12, 35) | — | — |  |
| CA | 18 (9.8, 39) | 0.88 | 0.49, 1.59 | 0.7 |
| CDCA | 25 (13, 55) | 0.72 | 0.39, 1.31 | 0.3 |
| LB^1^ |  |  |  |  |
| Veh | 16 (8.1, 34) | — | — |  |
| CA | 9.0 (5.4, 25) | 1.55 | 0.86, 2.77 | 0.14 |
| CDCA | 7.5 (5.0, 34) | 1.40 | 0.78, 2.52 | 0.3 |
| ^1^Log-Rank Test p-value (Within LB): p=0.3  ^2^Log-Rank Test p-value (Within Ctrl): p=0.5 | | | | |
| Abbreviations: CI = Confidence Interval, HR = Hazard Ratio | | | | |

Supplementary Table 12. Log-rank test comparing hazard ratios (HR) of CA and CDCA treated infant pups’ latency to approach the Dam against their Veh counterparts within each bedding condition. Median latency along with the 95% confidence interval (CI) was calculated for all pups. HR, 95% CI of HR, and p-value was calculated for Veh vs CA and Veh vs CDCA.

|  | Descriptive Summary | Cox Model | | |
| --- | --- | --- | --- | --- |
| Characteristic | Median (95% CI) | HR | 95% CI | p-value |
| Ctrl, Female^1^ |  |  |  |  |
| Veh | 51 (43, —) | — | — |  |
| CA | 133 (38, —) | 0.63 | 0.25, 1.61 | 0.3 |
| CDCA | 77 (46, —) | 1.02 | 0.43, 2.42 | >0.9 |
| Ctrl, Male^2^ |  |  |  |  |
| Veh | 50 (30, —) | — | — |  |
| CA | 143 (46, —) | 0.57 | 0.25, 1.32 | 0.2 |
| CDCA | 64 (25, —) | 0.66 | 0.28, 1.57 | 0.3 |
| LB, Female^3^ |  |  |  |  |
| Veh | 294 (215, —) | — | — |  |
| CA | 225 (34, —) | 1.56 | 0.58, 4.20 | 0.4 |
| CDCA | 84 (43, —) | 2.23 | 0.87, 5.73 | 0.10 |
| LB, Male^4^ |  |  |  |  |
| Veh | 178 (44, —) | — | — |  |
| CA | 66 (36, —) | 1.93 | 0.80, 4.67 | 0.15 |
| CDCA | 56 (24, —) | 2.30 | 0.94, 5.64 | 0.069 |
| ^1^Log-Rank Test p-value (Within Ctrl, F): p=0.5  ^2^Log-Rank Test p-value (Within Ctrl, M): p=0.4  ^3^Log-Rank Test p-value (Within LB, F): p=0.2  ^4^Log-Rank Test p-value (Within LB, M): p=0.2 | | | | |
| Abbreviations: CI = Confidence Interval, HR = Hazard Ratio | | | | |

Supplemental Table 13. Log-rank test comparing hazard ratios (HR) of Female and Male Ctrl and LB exposed infant pups’ latency to approach the ventrum against their Veh treated counterparts. Median latency along with the 95% confidence interval (CI) was calculated for all pups. HR, 95% CI of HR, and p-value was calculated for Veh vs CA and Veh vs CDCA.

|  | Descriptive Summary | Cox Model | | |
| --- | --- | --- | --- | --- |
| Characteristic | Median (95% CI) | HR | 95% CI | p-value |
| Veh, Female^1^ |  |  |  |  |
| Ctrl | 51 (43, —) | — | — |  |
| LB | 294 (215, —) | 0.50 | 0.19, 1.33 | 0.2 |
| Veh, Male^2^ |  |  |  |  |
| Ctrl | 50 (30, —) | — | — |  |
| LB | 178 (44, —) | 0.34 | 0.13, 0.90 | 0.029 |
| CA, Female^3^ |  |  |  |  |
| Ctrl | 133 (38, —) | — | — |  |
| LB | 225 (34, —) | 1.20 | 0.46, 3.11 | 0.7 |
| CA, Male^4^ |  |  |  |  |
| Ctrl | 143 (46, —) | — | — |  |
| LB | 66 (36, —) | 1.33 | 0.59, 2.98 | 0.5 |
| CDCA, Female^5^ |  |  |  |  |
| Ctrl | 77 (46, —) | — | — |  |
| LB | 84 (43, —) | 0.91 | 0.39, 2.08 | 0.8 |
| CDCA, Male^6^ |  |  |  |  |
| Ctrl | 64 (25, —) | — | — |  |
| LB | 56 (24, —) | 1.19 | 0.51, 2.79 | 0.7 |
| ^1^Log-Rank Test p-value (Within Veh, F): p=0.2  ^2^Log-Rank Test p-value (Within Veh, M): p=0.023  ^3^Log-Rank Test p-value (Within CA, F): p=0.7  ^4^Log-Rank Test p-value (Within CA, M): p=0.5  ^5^Log-Rank Test p-value (Within CDCA, F): p=0.8  ^6^Log-Rank Test p-value (Within CDCA, M): p=0.7 | | | | |
| Abbreviations: CI = Confidence Interval, HR = Hazard Ratio | | | | |

Supplemental Table 14. Log-rank test comparing hazard ratios (HR) of Female and Male Veh, CA, and CDCA treated infant pups’ latency to approach the ventrum against their Ctrl counterparts. Median latency along with the 95% confidence interval (CI) was calculated for all pups. HR, 95% CI of HR, and p-value was calculated for Veh vs CA and Veh vs CDCA.

|  | Descriptive Summary | Cox Model | | |
| --- | --- | --- | --- | --- |
| Characteristic^1^ | Median (95% CI) | HR | 95% CI | p-value |
| Ctrl, Veh^1^ |  |  |  |  |
| M | 50 (30, —) | — | — |  |
| F | 51 (43, —) | 0.71 | 0.29, 1.72 | 0.4 |
| Ctrl, CA^2^ |  |  |  |  |
| M | 143 (46, —) | — | — |  |
| F | 133 (38, —) | 0.60 | 0.24, 1.50 | 0.3 |
| Ctrl, CDCA^3^ |  |  |  |  |
| M | 64 (25, —) | — | — |  |
| F | 77 (46, —) | 1.06 | 0.46, 2.47 | 0.9 |
| LB, Veh^4^ |  |  |  |  |
| M | 178 (44, —) | — | — |  |
| F | 294 (215, —) | 0.87 | 0.32, 2.33 | 0.8 |
| LB, CA^5^ |  |  |  |  |
| M | 66 (36, —) | — | — |  |
| F | 225 (34, —) | 0.54 | 0.22, 1.33 | 0.2 |
| LB, CDCA^6^ |  |  |  |  |
| M | 56 (24, —) | — | — |  |
| F | 84 (43, —) | 0.78 | 0.34, 1.79 | 0.6 |
| ^1^Log-Rank Test p-value (Within Ctrl, Veh): p=0.4  ^2^Log-Rank Test p-value (Within Ctrl, CA): p=0.3  ^3^Log-Rank Test p-value (Within Ctrl, CDCA): p=0.9  ^4^Log-Rank Test p-value (Within LB, Veh): p=0.8  ^5^Log-Rank Test p-value (Within LB, CA): p=0.2  ^6^Log-Rank Test p-value (Within LB, CDCA): p=0.6 | | | | |
| Abbreviations: CI = Confidence Interval, HR = Hazard Ratio | | | | |

Supplemental Table 15. Log-rank test comparing hazard ratios (HR) of Veh, CA, and CDCA treated Ctrl and LB infant Female (F) pups’ latency to approach the ventrum against their Male (M) counterparts. Median latency along with the 95% confidence interval (CI) was calculated for all pups. HR, 95% CI of HR, and p-value was calculated for Veh vs CA and Veh vs CDCA.

|  | Descriptive Summary | Cox Model | | |
| --- | --- | --- | --- | --- |
| Characteristic | Median (95% CI) | HR | 95% CI | p-value |
| Ctrl, Female^1^ |  |  |  |  |
| Veh_oral | 24 (15, —) | — | — |  |
| CA | 12 (5.4, —) | 1.29 | 0.56, 3.00 | 0.5 |
| CDCA | 34 (13, —) | 1.01 | 0.44, 2.35 | >0.9 |
| Ctrl, Male^2^ |  |  |  |  |
| Veh_oral | 15 (11, —) | — | — |  |
| CA | 22 (14, —) | 0.52 | 0.22, 1.25 | 0.15 |
| CDCA | 24 (8.4, —) | 0.44 | 0.17, 1.10 | 0.079 |
| LB, Female^3^ |  |  |  |  |
| Veh_oral | 14 (7.4, —) | — | — |  |
| CA | 9.3 (2.6, —) | 1.35 | 0.57, 3.20 | 0.5 |
| CDCA | 6.4 (4.0, —) | 1.27 | 0.54, 2.97 | 0.6 |
| LB, Male^4^ |  |  |  |  |
| Veh_oral | 21 (7.9, —) | — | — |  |
| CA | 8.9 (5.3, —) | 1.79 | 0.79, 4.04 | 0.2 |
| CDCA | 8.7 (6.4, —) | 1.56 | 0.68, 3.56 | 0.3 |
| ^1^Log-Rank Test p-value (Within Ctrl, F): p=0.8  ^2^Log-Rank Test p-value (Within Ctrl, M): p=0.2  ^3^Log-Rank Test p-value (Within LB, F): p=0.8  ^4^Log-Rank Test p-value (Within LB, M): p=0.3 | | | | |
| Abbreviations: CI = Confidence Interval, HR = Hazard Ratio | | | | |

Supplemental Table 16. Log-rank test comparing hazard ratios (HR) of Female and Male Ctrl and LB exposed infant pups’ latency to approach the Dam against their Veh treated counterparts. Median latency along with the 95% confidence interval (CI) was calculated for all pups. HR, 95% CI of HR, and p-value was calculated for Veh vs CA and Veh vs CDCA.

|  | Descriptive Summary | Cox Model | | |
| --- | --- | --- | --- | --- |
| Characteristic | Median (95% CI) | HR | 95% CI | p-value |
| Veh, Female^1^ |  |  |  |  |
| Ctrl | 24 (15, —) | — | — |  |
| LB | 14 (7.4, —) | 1.28 | 0.54, 3.05 | 0.6 |
| Veh, Male^2^ |  |  |  |  |
| Ctrl | 15 (11, —) | — | — |  |
| LB | 21 (7.9, —) | 0.56 | 0.23, 1.39 | 0.2 |
| CA, Female^3^ |  |  |  |  |
| Ctrl | 12 (5.4, —) | — | — |  |
| LB | 9.3 (2.6, —) | 1.42 | 0.61, 3.29 | 0.4 |
| CA, Male^4^ |  |  |  |  |
| Ctrl | 22 (14, —) | — | — |  |
| LB | 8.9 (5.3, —) | 2.37 | 1.00, 5.59 | 0.050 |
| CDCA, Female^5^ |  |  |  |  |
| Ctrl | 34 (13, —) | — | — |  |
| LB | 6.4 (4.0, —) | 1.37 | 0.60, 3.15 | 0.5 |
| CDCA, Male^6^ |  |  |  |  |
| Ctrl | 24 (8.4, —) | — | — |  |
| LB | 8.7 (6.4, —) | 1.66 | 0.71, 3.89 | 0.2 |
| ^1^Log-Rank Test p-value (Within Veh, F): p=0.6  ^2^Log-Rank Test p-value (Within Veh, M): p=0.2  ^3^Log-Rank Test p-value (Within CA, F): p=0.4  ^4^Log-Rank Test p-value (Within CA, M): p=0.043  ^5^Log-Rank Test p-value (Within CDCA, F): p=0.5  ^6^Log-Rank Test p-value (Within CDCA, M): p=0.2 | | | | |
| Abbreviations: CI = Confidence Interval, HR = Hazard Ratio | | | | |

Supplemental Table 17. Log-rank test comparing hazard ratios (HR) of Female and Male Veh, CA, and CDCA treated infant pups’ latency to approach the ventrum against their Ctrl counterparts. Median latency along with the 95% confidence interval (CI) was calculated for all pups. HR, 95% CI of HR, and p-value was calculated for Veh vs CA and Veh vs CDCA.

|  | Descriptive Summary | Cox Model | | |
| --- | --- | --- | --- | --- |
| Characteristic | Median (95% CI) | HR | 95% CI | p-value |
| Ctrl, Veh |  |  |  |  |
| M | 15 (11, —) | — | — |  |
| F | 24 (15, —) | 0.40 | 0.15, 1.05 | 0.064 |
| Ctrl, CA^2^ |  |  |  |  |
| M | 22 (14, —) | — | — |  |
| F | 12 (5.4, —) | 1.07 | 0.47, 2.45 | 0.9 |
| Ctrl, CDCA^3^ |  |  |  |  |
| M | 24 (8.4, —) | — | — |  |
| F | 34 (13, —) | 1.09 | 0.47, 2.55 | 0.8 |
| LB, Veh^5^ |  |  |  |  |
| M | 21 (7.9, —) | — | — |  |
| F | 14 (7.4, —) | 1.07 | 0.47, 2.45 | 0.9 |
| LB, CA^5^ |  |  |  |  |
| M | 8.9 (5.3, —) | — | — |  |
| F | 9.3 (2.6, —) | 0.86 | 0.37, 1.98 | 0.7 |
| LB, CDCA^6^ |  |  |  |  |
| M | 8.7 (6.4, —) | — | — |  |
| F | 6.4 (4.0, —) | 0.88 | 0.38, 2.02 | 0.8 |
| ^1^Log-Rank Test p-value (Within Ctrl, Vehl): p=0.056  ^2^Log-Rank Test p-value (Within Ctrl, CA): p=0.9  ^3^Log-Rank Test p-value (Within Ctrl, CDCA): p=0.8  ^4^Log-Rank Test p-value (Within LB, Veh_oral): p=0.9  ^5^Log-Rank Test p-value (Within LB, CA): p=0.7  ^6^Log-Rank Test p-value (Within LB, CDCA): p=0.8 | | | | |
| Abbreviations: CI = Confidence Interval, HR = Hazard Ratio | | | | |

Supplemental Table 18. Log-rank test comparing hazard ratios (HR) of Veh, CA, and CDCA treated Ctrl and LB infant Female (F) pups’ latency to approach the Dam against their Male (M) counterparts. Median latency along with the 95% confidence interval (CI) was calculated for all pups. HR, 95% CI of HR, and p-value was calculated for M v F.
