## Supplementary Figures for "Bile acid signaling as a therapeutically tractable pathway linking early caregiving adversity to social behavior"

**Contents**

**Supplementary Figure 1.** Early social but not non-social stress perturbs infant-Dam social interaction.

**Supplementary Figure 2.** Differential effects of early social and non-social adversity on pup development and locomotion during infancy and adolescence.

**Supplementary Figure 3.** Differential effects of early social and non-social adversity on infant pup development and locomotion are sex specific.

**Supplementary Figure 4.** Differential effects of early social and non-social adversity on adolescent pup development and locomotion are sex-specific.

**Supplementary Figure 5.** Early social but not non-social stress affects infant social behavior in a sex-specific manner.

**Supplementary Figure 6.** Early social but not non-social stress perturbs adolescent social behavior in a sex-specific manner.

**Supplementary Figure 7.** Effects of social and nonsocial ELS on the transcriptomic profile of the infant female BLA.

**Supplementary Figure 8.** Effects of social and nonsocial ELS on the transcriptomic profile of the infant male BLA.

**Supplementary Figure 9.** Effects of social and nonsocial ELS on the transcriptomic profile of the adolescent female BLA.

**Supplementary Figure 10.** Effects of social and nonsocial ELS on the transcriptomic profile of the adolescent male BLA.

**Supplementary Figure 11.** Social and non-social ELS differentially alter infant female blood serum metabolome following MSA.

**Supplementary Figure 12.** Social and non-social ELS differentially alter infant male blood serum metabolome following MSA.

**Supplementary Figure 13.** Social and non-social ELS alter bile acid metabolism following 2-CSP in female adolescent pups.

**Supplementary Figure 14.** Social and non-social ELS disrupt alanine, glycogen, and pyruvate metabolism following 2-CSP in male adolescent pups.

**Supplementary Figure 15.** LB exposure does not perturb broad infant approach to Dam.

**Supplementary Figure 16.** Bile acid supplementation does not alter infant development nor locomotor activity during MSA.

**Supplementary Figure 17.** LB exposure does not perturb infant interaction with the ventrum in a sex-specific manner.

**Supplementary Figure 18.** LB exposure does not produce lasting changes in social behavior in the 2-CSP.

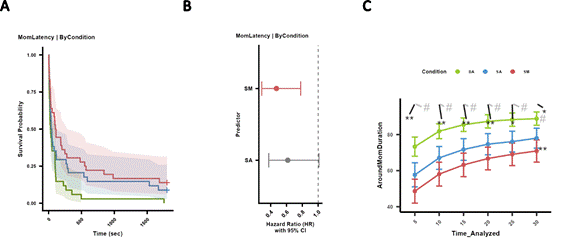

Supplementary Figure 1. Early social but not non-social stress perturbs infant-Dam social interaction. (A) Kaplan-Meier survival curves of BA, SA, and SM treated infant pups’ latency to approach the Dam (i.e., mom). Survival probability (y-axis) of approaching the ventrum is graphed for each treatment as a function of Time (sec). (B) Cox-proportional hazard and 95% confidence interval (CI) of latency to approach the Dam by SA and SM treated relative to BA treated infant pup. Red indicates significantly different and gray indicates no significant difference. (C) Line graph of proportion of total duration (%) spent around the Dam by the BA, SA, and SM treated infant pups is plotted for each 5 min time bin. Area under the curve of proportion of total duration spent around the Dam by BA, SA, and SM treated pups is also compared across each comparison and plotted to the right of the 30 min time bin value. Green is BA, blue is SA, and red is SM. # indicates p < 0.1, * indicates p < 0.05, ** indicates p < 0.01, and *** indicates p < 0.001. Additionally, light gray * or # reflect comparison between BA and SA, black * or # reflect comparison between BA and SM, and dark gray * or # reflect comparison between SA and SM.

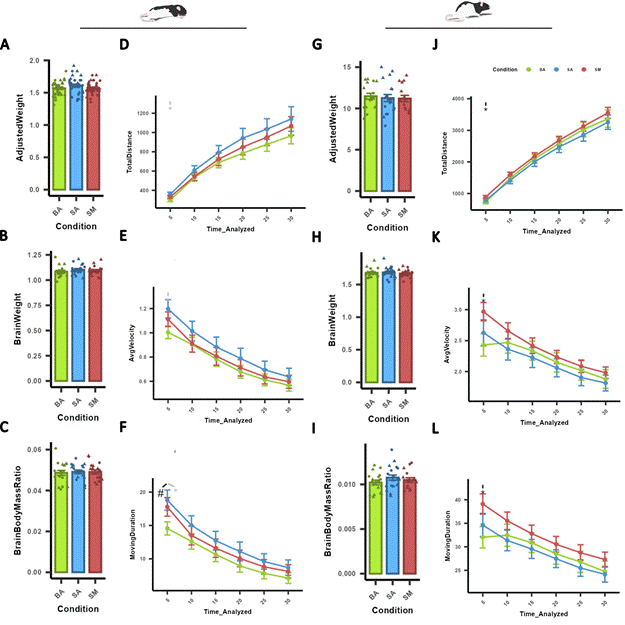

Supplementary Figure 2. Differential effects of early social and non-social adversity on pup development and locomotion during infancy and adolescence. Bar plots of infant (A-C) and adolescent (G-I) pup weight relative to P8 timepoint (AdjustedWeight), brain weight in g (BrainWeight), and brain to body mass ratio (BrainBodyMassRatio). Line graphs of total distance traveled (cm), average velocity (cm/s), and total time spent moving (MovingDuration, s) are plotted for each 5 min time bin during the infant MSA (D-F) and adolescent 2-CSP (J-L). Area under the curve of each metric was also compared across each comparison and plotted to the right of the 30 min time bin value. Green is BA, blue is SA, and red is SM. # indicates p < 0.1, * indicates p < 0.05, ** indicates p < 0.01, and *** indicates p < 0.001. Additionally, light gray * or # reflect comparison between BA and SA, black * or # reflect comparison between BA and SM, and dark gray * or # reflect comparison between SA and SM.

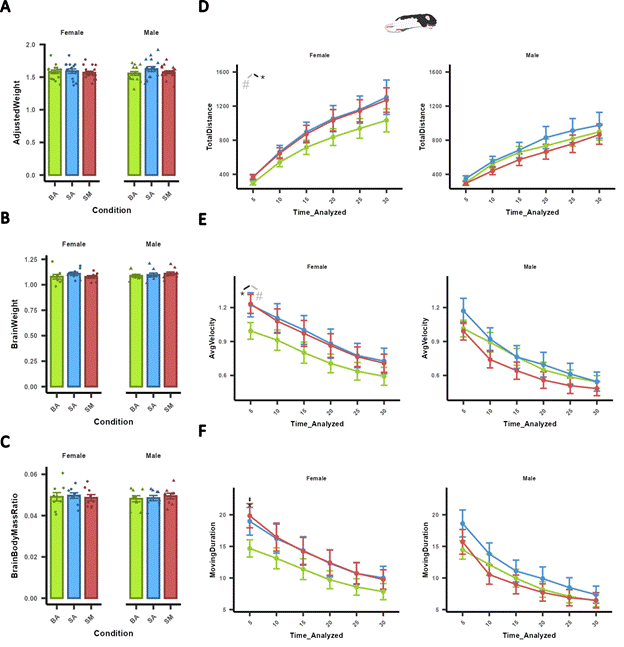

Supplementary Figure 3. Differential effects of early social and non-social adversity on infant pup development and locomotion are sex specific. Bar plots of infant (A-C) pup weight relative to P8 timepoint (AdjustedWeight), brain weight in g (BrainWeight), and brain to body mass ratio (BrainBodyMassRatio). Line graphs of total distance traveled (cm), average velocity (cm/s), and total time spent moving (MovingDuration, s) are plotted for each 5 min time bin by sex during the infant MSA (D-F). Area under the curve of each metric was also compared across each comparison and plotted to the right of the 30 min time bin value. Green is BA, blue is SA, and red is SM. # indicates p < 0.1, * indicates p < 0.05, ** indicates p < 0.01, and *** indicates p < 0.001. Additionally, light gray * or # reflect comparison between BA and SA, black * or # reflect comparison between BA and SM, and dark gray * or # reflect comparison between SA and SM.

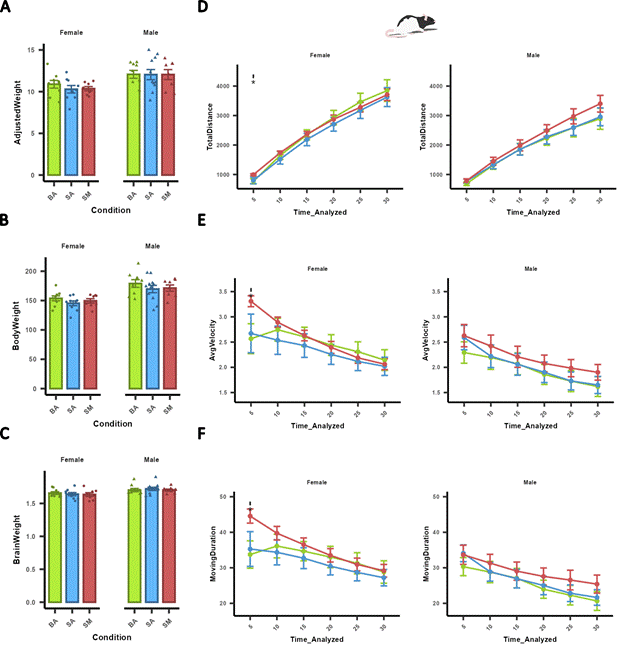

Supplementary Figure 4. Differential effects of early social and non-social adversity on adolescent pup development and locomotion are sex-specific. Bar plots of adolescent (A-C) pup weight relative to P8 timepoint (AdjustedWeight), brain weight in g (BrainWeight), and brain to body mass ratio (BrainBodyMassRatio). Line graphs of total distance traveled (cm), average velocity (cm/s), and total time spent moving (MovingDuration, s) are plotted for each 5 min time bin by sex during the adolescent 2-CSP (D-F). Area under the curve of each metric was also compared across each comparison and plotted to the right of the 30 min time bin value. Green is BA, blue is SA, and red is SM. # indicates p < 0.1, * indicates p < 0.05, ** indicates p < 0.01, and *** indicates p < 0.001. Additionally, light gray * or # reflect comparison between BA and SA, black * or # reflect comparison between BA and SM, and dark gray * or # reflect comparison between SA and SM.

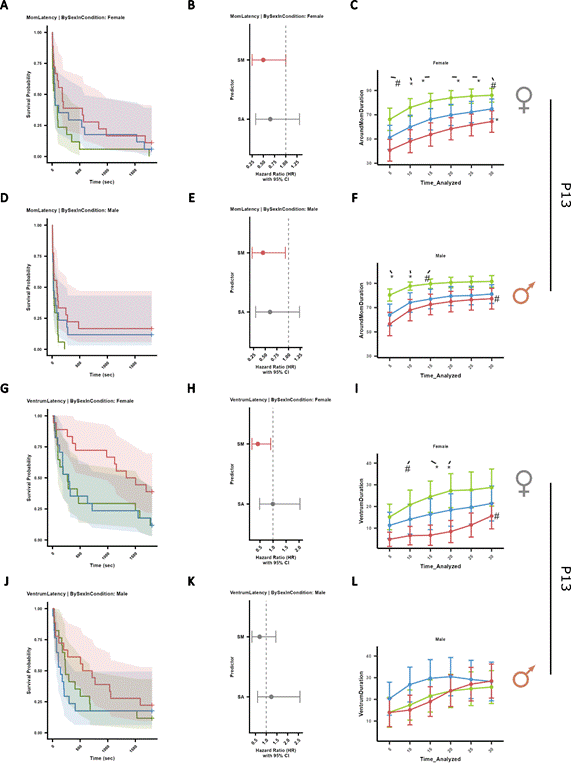

Supplementary Figure 5. Early social but not non-social stress affects infant social behavior in a sex-specific manner. (A, D) Kaplan-Meier survival curves of BA, SA, and SM treated female (A) and male (D) infant pups’ latency to approach the Dam. Survival probability (y-axis) of approaching the Dam is graphed for each treatment as a function of Time (sec). (B, E) Cox-proportional hazard and 95% confidence interval (CI) of latency to approach the Dam by female (B) and male (E) SA and SM treated relative to BA treated infant pup. Red indicates significantly different and gray indicates no significant difference. (C, F) Line graph of proportion of total duration (%) spent around the Dam by female (C) and male (F) BA, SA, and SM treated infant pups is plotted for each 5 min time bin. Area under the curve of proportion of total duration spent around the Dam by BA, SA, and SM treated pups is also compared across each comparison and plotted to the right of the 30 min time bin value. (G, J) Kaplan-Meier survival curves of BA, SA, and SM treated female (G) and male (J) infant pups’ latency to approach the ventrum. Survival probability (y-axis) of approaching the ventrum is graphed for each treatment as a function of Time (sec). (H, K) Cox-proportional hazard and 95% confidence interval (CI) of latency to approach the Dam by female (H) and male (K) SA and SM treated relative to BA treated infant pup. Red indicates significantly different and gray indicates no significant difference. (I, L) Line graph of proportion of total duration (%) spent around the Dam by female (I) and male (L) BA, SA, and SM treated infant pups is plotted for each 5 min time bin. Area under the curve of proportion of total duration spent around the Dam by BA, SA, and SM treated pups is also compared across each comparison and plotted to the right of the 30 min time bin value. Green is BA, blue is SA, and red is SM. # indicates p < 0.1, * indicates p < 0.05, ** indicates p < 0.01, and *** indicates p < 0.001. Additionally, light gray * or # reflect comparison between BA and SA, black * or # reflect comparison between BA and SM, and dark gray * or # reflect comparison between SA and SM.

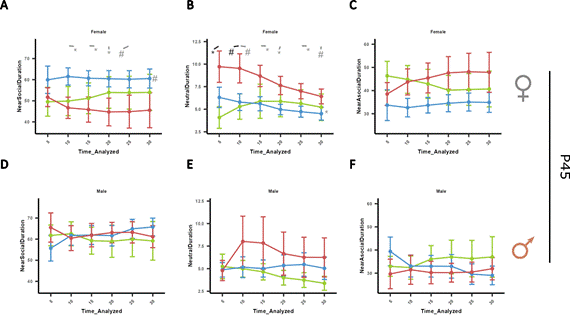

Supplementary Figure 6. Early social but not non-social stress perturbs adolescent social behavior in a sex-specific manner. (A-F) Line graph of proportion of total duration (%) spent near the social stim (NearSocial, A & D), neutral (middle, B &E), and near the empty (NearAsocial, C & F) side of the 2-CSP apparatus are plotted within each 5 min time bin for female (A-C) and male (D-F) adolescent pups. Area under the curve of proportion of total duration spent around or within a region of interest is compared across each comparison (BA vs SA, BA vs SM, SM vs SA) and plotted to the right of the 30 min time bin value. Green is BA, blue is SA, and red is SM. # indicates p < 0.1, * indicates p < 0.05, ** indicates p < 0.01, and *** indicates p < 0.001. Additionally, light gray * or # reflect comparison between BA and SA, black * or # reflect comparison between BA and SM, and dark gray * or # reflect comparison between SA and SM.

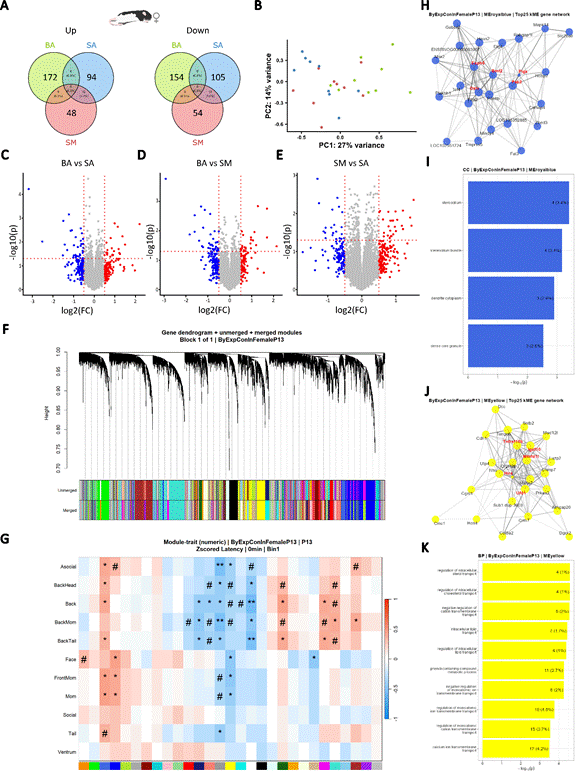

Supplementary Figure 7. Effects of social and nonsocial ELS on the transcriptomic profile of the infant female BLA. (A) Venn diagram of up (left) and down (right) regulated genes. Values represent number of genes up or down regulated in treatment conditions relative to BA. (B) PCA of infant female BA, SA, and SM BLA’s transcriptomic profile. (C, D, E) Volcano plots highlighting significantly up- and down-regulated genes for each pair-wise comparison. Left to right, BA vs SA, BA vs SM, and SM vs SA. -log10 of the pvalue is plotted on the y-axis, and log2 of the fold change (FC) between both conditions is plotted on the x-axis. Horizontal red dashed line drawn at Veritical red dashed lines drawn at -0.25 and -.25. Red and blue indicate a FC of ± 0.5, respectively. Gray indicates a FC between -0.5 and 0.5. (F) Weighted Gene CoExpression Network Analysis of infant (P13) samples reveals that genes in the infant female BLA cluster into distinct modules. Each branch in the dendrogram represents a gene. Each gene is assigned to a cluster based on its distance to other genes, resulting in unmerged modules. Modules that are 75% correlated are merged. (G) Heatmap of correlations between merged modules and latency to approach infant pup data reveals novel module-trait relationships. Red and blue indicate positive and negative correlation, respectively. # indicates p < 0.1, * indicates p < 0.05, ** indicates p < 0.01, and *** indicates p < 0.001. (H, J) Network visualization of top 25 hub-genes in green (H) and turquoise (J) module by kME graphed using Fruchterman-Reingold layout. Red text highlights the five hub-genes with the highest average kME. (I, K) Overrepresentation Analysis of genes in the royal blue (I) and yellow (K) modules reveal stereocilia, dendrite, and sterol plus cholesterol transport as key regulators of social behavior in infant female BLA.

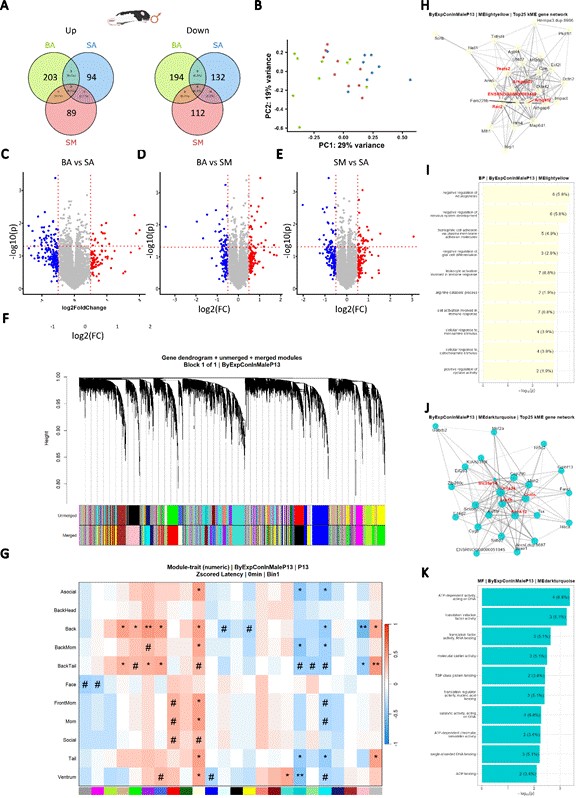

Supplementary Figure 8. Effects of social and nonsocial ELS on the transcriptomic profile of the infant male BLA. (A) Venn diagram of up (left) and down (right) regulated genes. Values represent number of genes up or down regulated in treatment conditions relative to BA. (B) PCA of infant male BA, SA, and SM BLA’s transcriptomic profile. (C, D, E) Volcano plots highlighting significantly up- and down-regulated genes for each pair-wise comparison. Left to right, BA vs SA, BA vs SM, and SM vs SA. -log10 of the pvalue is plotted on the y-axis, and log2 of the fold change (FC) between both conditions is plotted on the x-axis. Horizontal red dashed line drawn at Veritical red dashed lines drawn at -0.25 and -.25. Red and blue indicate a FC of ± 0.5, respectively. Gray indicates a FC between -0.5 and 0.5. (F) Weighted Gene CoExpression Network Analysis of infant (P13) samples reveals that genes in the infant male BLA cluster into distinct modules. Each branch in the dendrogram represents a gene. Each gene is assigned to a cluster based on its distance to other genes, resulting in unmerged modules. Modules that are 75% correlated are merged. (G) Heatmap of correlations between merged modules and latency to approach infant pup data reveals novel module-trait relationships. Red and blue indicate positive and negative correlation, respectively. # indicates p < 0.1, * indicates p < 0.05, ** indicates p < 0.01, and *** indicates p < 0.001. (H, J) Network visualization of top 25 hub-genes in green (H) and turquoise (J) module by kME graphed using Fruchterman-Reingold layout. Red text highlights the five hub-genes with the highest average kME. (I, K) Overrepresentation Analysis of genes in the light yellow (I) and cyan (K) modules reveal negative regulation of neurogenesis and nervous system development plus translation as key regulators of social behavior in infant male BLA.

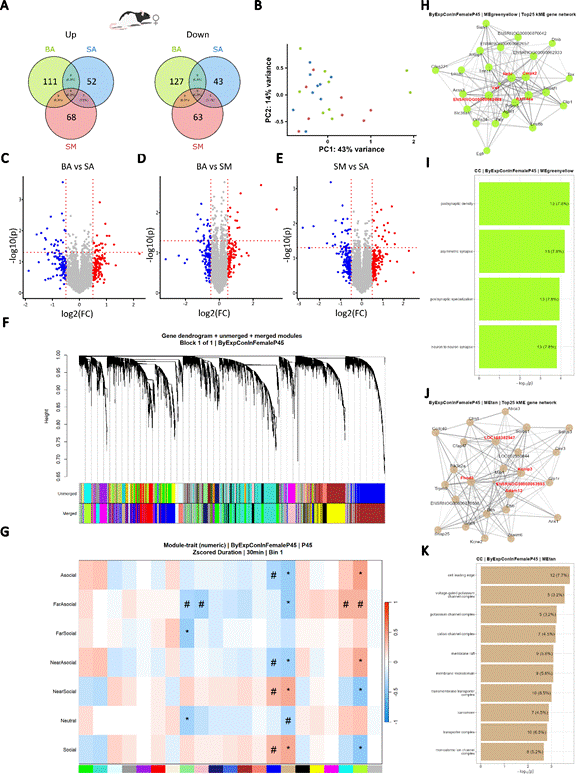

Supplemental Figure 9. Effects of social and nonsocial ELS on the transcriptomic profile of the adolescent female BLA. (A) Venn diagram of up (left) and down (right) regulated genes. Values represent number of genes up or down regulated in treatment conditions relative to BA. (B) PCA of adolescent female BA, SA, and SM BLA’s transcriptomic profile. (C, D, E) Volcano plots highlighting significantly up- and down-regulated genes for each pair-wise comparison. Left to right, BA vs SA, BA vs SM, and SM vs SA. -log10 of the pvalue is plotted on the y-axis, and log2 of the fold change (FC) between both conditions is plotted on the x-axis. Horizontal red dashed line drawn at Veritical red dashed lines drawn at -0.25 and -.25. Red and blue indicate a FC of ± 0.5, respectively. Gray indicates a FC between -0.5 and 0.5. (F) Weighted Gene CoExpression Network Analysis of adolescent (P45) samples reveals that genes in the adolescent female BLA cluster into distinct modules. Each branch in the dendrogram represents a gene. Each gene is assigned to a cluster based on its distance to other genes, resulting in unmerged modules. Modules that are 75% correlated are merged. (G) Heatmap of correlations between merged modules and duration pup data reveals novel module-trait relationships. Red and blue indicate positive and negative correlation, respectively. # indicates p < 0.1, * indicates p < 0.05, ** indicates p < 0.01, and *** indicates p < 0.001. (H, J) Network visualization of top 25 hub-genes in green (H) and turquoise (J) module by kME graphed using Fruchterman-Reingold layout. Red text highlights the five hub-genes with the highest average kME. (I, K) Overrepresentation Analysis of genes in the green yellow (I) and tan (K) modules reveal postsynaptic density, synapse, and potassium channels as key regulators of social behavior in adolescent female BLA.

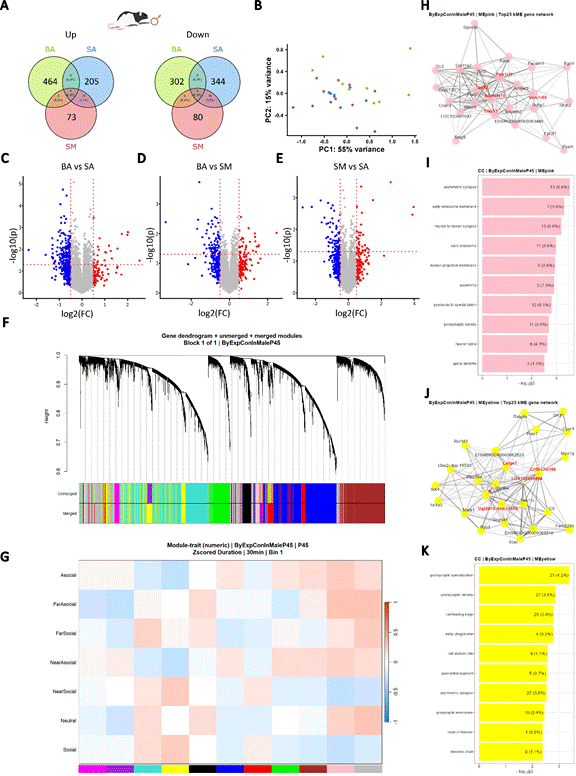

Supplemental Figure 10. Effects of social and nonsocial ELS on the transcriptomic profile of the adolescent male BLA. (A) Venn diagram of up (left) and down (right) regulated genes. Values represent number of genes up or down regulated in treatment conditions relative to BA. (B) PCA of adolescent male BA, SA, and SM BLA’s transcriptomic profile. (C, D, E) Volcano plots highlighting significantly up- and down-regulated genes for each pair-wise comparison. Left to right, BA vs SA, BA vs SM, and SM vs SA. -log10 of the pvalue is plotted on the y-axis, and log2 of the fold change (FC) between both conditions is plotted on the x-axis. Horizontal red dashed line drawn at Veritical red dashed lines drawn at -0.25 and -.25. Red and blue indicate a FC of ± 0.5, respectively. Gray indicates a FC between -0.5 and 0.5. (F) Weighted Gene CoExpression Network Analysis of adolescent (P45) samples reveals that genes in the adolescent female BLA cluster into distinct modules. Each branch in the dendrogram represents a gene. Each gene is assigned to a cluster based on its distance to other genes, resulting in unmerged modules. Modules that are 75% correlated are merged. (G) Heatmap of correlations between merged modules and duration pup data reveals novel module-trait relationships. Red and blue indicate positive and negative correlation, respectively. # indicates p < 0.1, * indicates p < 0.05, ** indicates p < 0.01, and *** indicates p < 0.001. (H, J) Network visualization of top 25 hub-genes in green (H) and turquoise (J) module by kME graphed using Fruchterman-Reingold layout. Red text highlights the five hub-genes with the highest average kME. (I, K) Overrepresentation Analysis of genes in the green yellow (I) and tan (K) modules reveal synapse, endosomes, and postsynaptic density as key regulators of social behavior in adolescent male BLA.

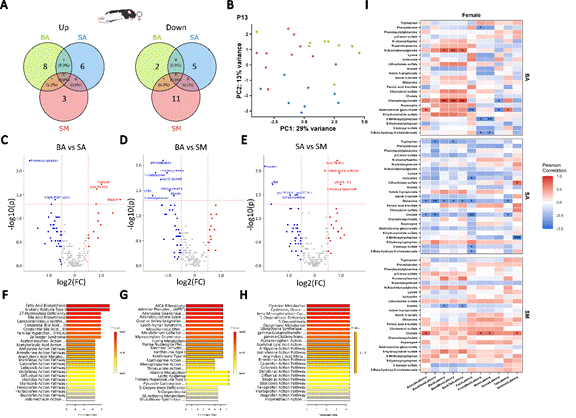

Supplementary Figure 11. Social and non-social ELS differentially alter infant female blood serum metabolome following MSA. (A) Venn diagram of up (left) and down (right) regulated metabolites in infant blood serum. Values represent number of metabolites up or down regulated in each condition. (B) PCA of infant female BA, SA, and SM’s blood serum metabolome. (C, D, E) Volcano plots highlighting significantly up- and down-regulated metabolites for each pair-wise comparison. Left to right, BA vs SA, BA vs SM, and SM vs SA. -log10 of the pvalue is plotted on the y-axis, and log2 of the fold change (FC) between both conditions is plotted on the x-axis. Horizontal red dashed line drawn at Veritical red dashed lines drawn at -0.25 and -.25. Red and blue indicate a FC of ± 0.5, respectively. Gray indicates a FC between -0.5 and 0.5. Labeled metabolites indicate a significant (p < 0.05) FC of ± 0.5. (F, G, H) Overrepresentation Analysis of all significantly different metabolites within BA vs SA (F), BA vs SM (G), and SA vs SM (H) does not reveal significantly overrepresented pathways. (I) Heatmap of metabolite-latency correlations reveal treatment dependent relationships. Red and blue indicate positive and negative correlation, respectively. * indicates p < 0.05, ** indicates p < 0.01, and *** indicates p < 0.001.

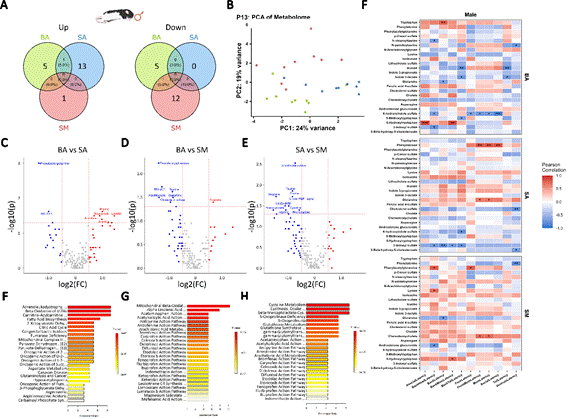

Supplementary Figure 12. Social and non-social ELS differentially alter infant male blood serum metabolome following MSA. (A) Venn diagram of up (left) and down (right) regulated metabolites in infant blood serum. Values represent number of metabolites up or down regulated in each condition. (B) PCA of infant female BA, SA, and SM’s blood serum metabolome. (C, D, E) Volcano plots highlighting significantly up- and down-regulated metabolites for each pair-wise comparison. Left to right, BA vs SA, BA vs SM, and SM vs SA. -log10 of the pvalue is plotted on the y-axis, and log2 of the fold change (FC) between both conditions is plotted on the x-axis. Horizontal red dashed line drawn at Veritical red dashed lines drawn at -0.25 and -.25. Red and blue indicate a FC of ± 0.5, respectively. Gray indicates a FC between -0.5 and 0.5. Labeled metabolites indicate a significant (p < 0.05) FC of ± 0.5. (F, G, H) Overrepresentation Analysis of all significantly different metabolites within BA vs SA (F), BA vs SM (G), and SA vs SM (H) does not reveal significantly overrepresented pathways. (I) Heatmap of metabolite-latency correlations reveal treatment dependent relationships. Red and blue indicate positive and negative correlation, respectively. * indicates p < 0.05, ** indicates p < 0.01, and *** indicates p < 0.001.

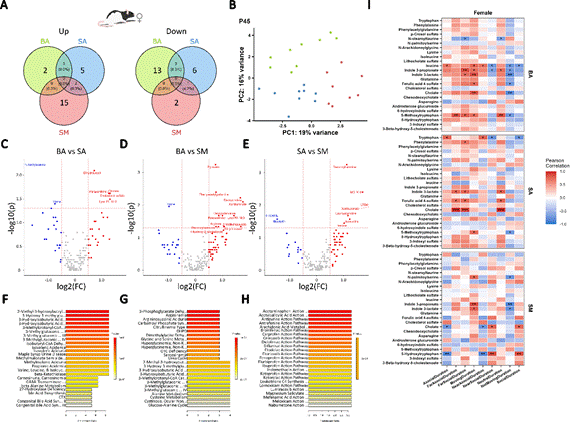

Supplemental Figure 13. Social and non-social ELS alter bile acid metabolism following 2-CSP in female adolescent pups. (A) Venn diagram of up (left) and down (right) regulated metabolites in infant blood serum. Values represent number of metabolites up or down regulated in each condition. (B) PCA of infant female BA, SA, and SM’s blood serum metabolome. (C, D, E) Volcano plots highlighting significantly up- and down-regulated metabolites for each pair-wise comparison. Left to right, BA vs SA, BA vs SM, and SM vs SA. -log10 of the pvalue is plotted on the y-axis, and log2 of the fold change (FC) between both conditions is plotted on the x-axis. Horizontal red dashed line drawn at Veritical red dashed lines drawn at -0.25 and -.25. Red and blue indicate a FC of ± 0.5, respectively. Gray indicates a FC between -0.5 and 0.5. Labeled metabolites indicate a significant (p < 0.05) FC of ± 0.5. (F, G, H) Overrepresentation Analysis of all significantly different metabolites within BA vs SA (F), BA vs SM (G), and SA vs SM (H) does not reveal significantly overrepresented pathways. (I) Heatmap of metabolite-duration correlations reveal bile acid metabolism disruption. Red and blue indicate positive and negative correlation, respectively. * indicates p < 0.05, ** indicates p < 0.01, and *** indicates p < 0.001.

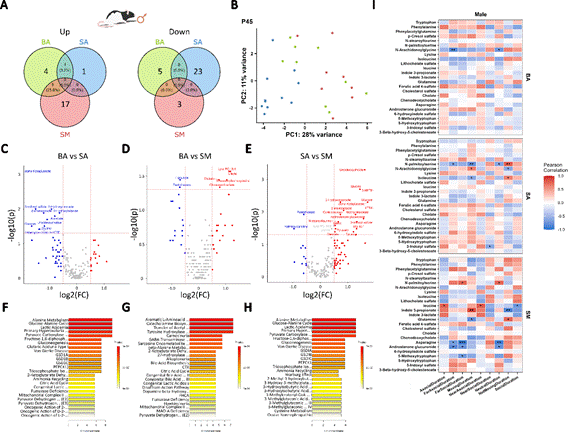

Supplemental Figure 14. Social and non-social ELS disrupt alanine, glycogen, and pyruvate metabolism following 2-CSP in male adolescent pups. (A) Venn diagram of up (left) and down (right) regulated metabolites in infant blood serum. Values represent number of metabolites up or down regulated in each condition. (B) PCA of infant female BA, SA, and SM’s blood serum metabolome. (C, D, E) Volcano plots highlighting significantly up- and down-regulated metabolites for each pair-wise comparison. Left to right, BA vs SA, BA vs SM, and SM vs SA. -log10 of the pvalue is plotted on the y-axis, and log2 of the fold change (FC) between both conditions is plotted on the x-axis. Horizontal red dashed line drawn at Veritical red dashed lines drawn at -0.25 and -.25. Red and blue indicate a FC of ± 0.5, respectively. Gray indicates a FC between -0.5 and 0.5. Labeled metabolites indicate a significant (p < 0.05) FC of ± 0.5. (F, G, H) Overrepresentation Analysis of all significantly different metabolites within BA vs SA (F), BA vs SM (G), and SA vs SM (H) reveal alanine, glycogen, and pyruvate metabolic pathways as overrepresented pathways. (I) Heatmap of metabolite-duration correlations reveal bile acid metabolism disruption. Red and blue indicate positive and negative correlation, respectively. * indicates p < 0.05, ** indicates p < 0.01, and *** indicates p < 0.001.

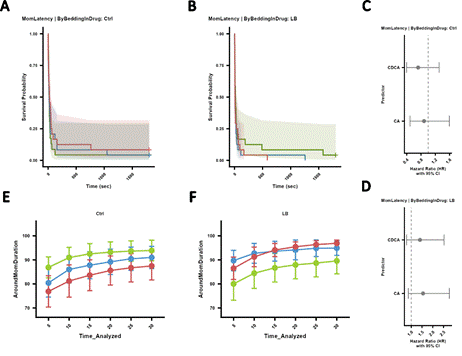

Supplementary Figure 15. LB exposure does not perturb broad infant approach to Dam. (A, B) Kaplan-Meier survival curves of Veh, CA, and CDCA treated infant Ctrl and LB pups’ latency to approach anywhere around the Dam. Survival probability (y-axis) of approaching the ventrum is graphed for each treatment as a function of Time (sec). (C, D) Cox-proportional hazard and 95% confidence interval (CI) of latency to approach the ventrum by CA and CDCA treated relative to Veh treated infant Ctrl and LB pups. Red indicates significantly different and gray indicates no significant difference. (E, F) Line graph of proportion of total duration (%) spent around the Dam by the Veh, CA, and CDCA treated infant pups is plotted for each 5 min time bin. Area under the curve of proportion of total duration spent around the Dam by Veh, CA, and CDCA treated Ctrl and LB pups is also compared across each comparison and plotted to the right of the 30 min time bin value. Green is Veh, blue is CA, and red is CDCA. # indicates p < 0.1, * indicates p < 0.05, ** indicates p < 0.01, and *** indicates p < 0.001. Additionally, light gray * or # reflect comparison between Veh and CA, black * or # reflect comparison between Veh and CDCA, and dark gray * or # reflect comparison between CA and CDCA.

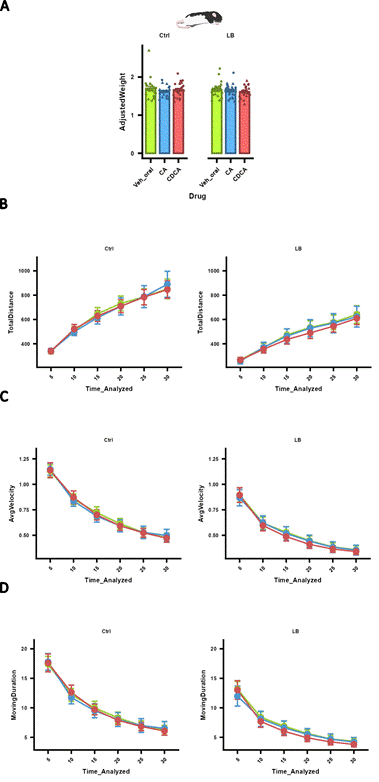

Supplemental Figure 16. Bile acid supplementation does not alter infant development nor locomotor activity during MSA. (A) Bar plot of infant pup weight relative to P8 timepoint (AdjustedWeight). (B-D) Line graphs of total distance traveled (cm), average velocity (cm/s), and total time spent moving (MovingDuration, s) are plotted for each 5 min time bin by bedding exposure (Ctrl and LB) during MSA. Area under the curve of each metric was also compared across each comparison (Veh vs CA, Veh vs CDCA, CDCA vs CA) and plotted to the right of the 30 min time bin value. Green is Veh, blue is CA, and red is CDCA. # indicates p < 0.1, * indicates p < 0.05, ** indicates p < 0.01, and *** indicates p < 0.001. Additionally, light gray * or # reflect comparison between Veh and CA, black * or # reflect comparison between Veh and CDCA, and dark gray * or # reflect comparison between CA and CDCA.

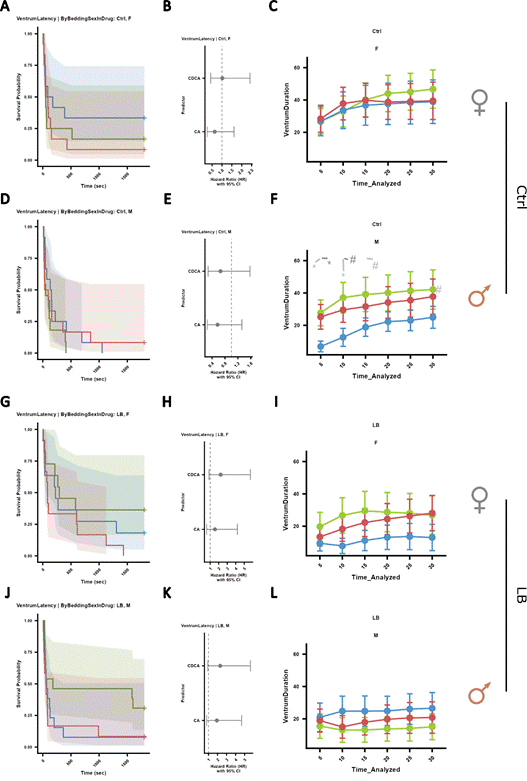

Supplementary Figure 17. LB exposure does not perturb infant interaction with the ventrum in a sex-specific manner. (A, D, G, J) Kaplan-Meier survival curves of Veh, CA, and CDCA treated infant Female and Male Ctrl and LB pups’ latency to approach the Dam’s ventrum. Survival probability (y-axis) of approaching the ventrum is graphed for each treatment as a function of Time (sec). (B, E, H, K) Cox-proportional hazard and 95% confidence interval (CI) of latency to approach the ventrum by CA and CDCA treated relative to Veh treated infant Ctrl and LB pups. Red indicates significantly different and gray indicates no significant difference. (C, F, I, L) Line graph of proportion of total duration (%) spent around the Dam by the Veh, CA, and CDCA treated infant pups is plotted for each 5 min time bin. Area under the curve of proportion of total duration spent around the Dam’s ventrum by Veh, CA, and CDCA treated Female and Male Ctrl and LB pups is also compared across each comparison and plotted to the right of the 30 min time bin value. Green is Veh, blue is CA, and red is CDCA. # indicates p < 0.1, * indicates p < 0.05, ** indicates p < 0.01, and *** indicates p < 0.001. Light gray * or # reflect comparison between Veh and CA, black * or # reflect comparison between Veh and CDCA, and dark gray * or # reflect comparison between CA and CDCA.

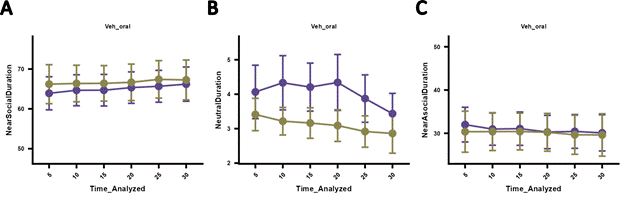

Supplementary Figure 18. LB exposure does not produce lasting changes in social behavior in the adolescent 2-CSP. (A-C) Line graph of proportion of total duration (%) spent near the social stim (NearSocial, A), neutral (middle, B), and near the empty (NearAsocial, C) side of the 2-CSP apparatus are plotted within each 5 min time bin for adolescent pups. Area under the curve of proportion of total duration spent around or within a region of interest is compared across the Ctrl vs LB comparison and plotted to the right of the 30 min time bin value. Purple is Ctrl and Khaki is LB. # indicates p < 0.1, * indicates p < 0.05, ** indicates p < 0.01, and *** indicates p < 0.001. Black * or # reflect comparison between Ctrl vs LB.
