## Supplementary Methods for "Bile acid signaling as a therapeutically tractable pathway linking early caregiving adversity to social behavior"

**Animals**

Female and male Long Evans rats were bred for these experiments at the Kennedy Krieger Institute. Litters of approximately 12 pups containing 6 males and 6 females were used for all experiments. Cross-fostering from untreated litters was performed as necessary. At most one male and one female were used from each litter per treatment group at each developmental time point to account for litter effects. All experimental animals were marked and weighed before undergoing adversity from P8 through P12, at weekly time points following adversity starting at P21 through P42, and before the infant and adolescent social behavior tests. Animals were marked to track individual pups during adversity treatment and then tagged with an RFID chip the day after infant social behavior to minimize pup and maternal stress and isoflurane exposure. Male and female pups were separated following weaning at P21. Blood and tissue were collected after behavior tests at P13 or P45 for deconstructed adversity experiments and between P45 to P50 for bile acid supplementation experiments.

**Deconstructed Adversity Model (DAM)**

In the DAM, 8 pups from a 12-pup litter were randomly assigned to one of three treatment groups from P8 to P12: beaker alone (control, BA), repeated shock alone (non-social adversity, SA), or shock in the presence of an anesthetized maternal-odor producing dam (social adversity, SM). BA pups were left alone on a heated pad for 90 minutes each day. SA pups were similarly left on a heated pad but additionally received 18 low-amplitude shocks delivered every 5 minutes each day. SM pups received the same shock protocol in the presence of a maternal-odor producing dam. The remaining four pups in each litter were left untouched to minimize maternal stress in the home cage. Prior to treatment exposure, animals were assigned, weighed, and marked. Two males and two females were randomly assigned to each treatment condition so that tissue from one male and one female could be collected following each social behavior test per condition.

**Scarcity Adversity Model via Limited Bedding (SAM-LB)**

In the SAM-LB model, litters underwent typical rearing until P8. From P8 to P12, bedding was either decreased to 100 cc (Limited Bedding, LB) or continued at 2000 cc (Control, Ctrl). LB cages were cleared of waste daily to prevent maternal nest building. Maternal and pup behaviors were recorded for 1 hour daily from P8 to P12 and hand-scored by highly trained raters using BORIS software.^44^ Total time spent, number of occurrences, proportion of time, and mean time per behavior were scored. The behaviors analyzed have previously been described by Opendak et al.^30^. Four additional behavioral indices were also quantified: maternal pup interaction (‘Mom Moves Pup(s)’), maternal self-directed activity (‘Mom Self-Activity’), maternal nursing (‘Mom Nursing Pup(s)’), and maternal maltreatment (‘Mom Maltreats Pup’). Control litters showing unusually high maltreatment and SAM-LB litters showing insufficient maltreatment were excluded from analyses.

**Bile Acid Supplementation**

To test the contribution of primary BAs to ELS-induced social behavior deficits, pups within each SAM-LB or Ctrl litter were orally administered 50 µL of either vehicle oil with 10% DMSO (Veh), 2.5 mg/kg cholic acid (CA), or 2.5 mg/kg chenodeoxycholic acid (CDCA) daily from P8 to P12. Animals were assigned, weighed, and marked at P8. At least one male and one female were assigned to each treatment condition per litter. Infant social behavior was assessed at P13-15 and adolescent social behavior at P45-50. Tissue was collected 60 minutes following P45-50 social behavior.

**Maternal Social Approach Test (Mom Pup Test)**

Infant pups (P13-15) underwent a 30-minute maternal social approach test with an unfamiliar urethane-anesthetized maternal-odor producing dam between 6 AM and 6 PM ET. Prior to testing, the dam was placed on her side on one end of a polypropylene cage containing a thin layer of sterilized corncob bedding to allow ventrum access. A single pup was placed on the opposing end and allowed to freely interact with the anesthetized dam for 30 minutes. Latency to approach, time spent near, and number of visits to the dam’s ventrum, face, tail, back, back of head, and rear zones were tracked, along with time spent in the away-from-dam zone, time spent moving and not moving, distance traveled, and average velocity. Dam placement was counterbalanced across litters. Pups designated for euthanasia in DAM experiments were collected without anesthesia 60 minutes after the test; pups in bile acid supplementation experiments were returned to the home cage after testing.

**2-Choice Social Preference Test (Pup Pup Test)**

Adolescent pups (P45-50) underwent a 30-minute two-choice social preference test (2-CSP) with an unfamiliar age- and sex-matched pup between 6 AM and 6 PM ET. A three-chamber rectangular polypropylene apparatus was used, consisting of a large middle chamber (16 in L × 8 in W × 10 in H) and two smaller chambers (6 in L × 8 in W × 10 in H) separated from the middle by transparent perforated ¼-inch propylene walls to allow odor flow. After a 5-minute habituation period in the middle chamber, a novel social stimulus pup was placed on one side. The apparatus was cleaned with 70% ethanol followed by ddH₂O between subjects, with a minimum of 10 minutes between pups. The side containing the social stimulus was alternated across subjects. Time spent near the social side, in the neutral middle zone, and near the asocial side were tracked in Ethovision v17,^45^ along with locomotor measures. All pups were collected without isoflurane anesthesia 60 minutes after testing.

**Blood Serum and Brain Tissue Collection**

Following social behavior tests, pups were singly housed in fresh corncob bedding for 60 minutes prior to euthanasia, a time point selected based on prior work showing peak cFos expression at this interval. Whole brains were immediately weighed, and flash frozen with dry ice following trunk blood collection (< 30 sec). Trunk blood was kept on ice for 3-6 minutes before centrifugation at 3500 × g for 15 minutes at 2°C to isolate serum. Whole brain and blood serum samples were stored at -80°C.

For BLA dissection, flash-frozen brains were equilibrated on a cryostat at -20°C for 1 hour. The infant and adolescent basolateral amygdala was bilaterally harvested using stereotaxic coordinates from Khazipov et al.^46^. A 1 mm deep tissue punch was collected using a 20G Luer adapter and syringe into 1.5 mL PCR-clean tubes containing 15 µL ice-cold RNAprotect buffer (Qiagen, #76104) and stored at -80°C until mRNA isolation. Tissue sections before, within, and after the punched area were plated on charged slides for Nissl staining to verify bilateral punch accuracy.

**Blood Serum Metabolomics**

Metabolites were extracted from 100 µL of blood serum using 900 µL of 70% ice-cold methanol containing 0.5% 1 N HCl and pre-spiked heavy isotope labeled internal standards (Laurate-D23, Octanoate-D15, Caffeine-13C3, Citrate-D4, Lauroylcarnitine-D3, Glutamate-D3, Chenodeoxycholate-D5, Tryptophan-D5, Tyrosine-D2, and 18:1-d7 Lyso PC). Following protein precipitation, samples were vortexed for 5 min at 50 Hz using a TissueLyser LT (Qiagen), centrifuged at 13,000 rpm for 20 min at 4°C, and the resulting supernatant was evaporated to dryness using a vacuum concentrator (Thermo Scientific Savant SpeedVac SPD120P2). The dried residue, containing both endogenous metabolites and internal standards, was reconstituted in 150 µL of 50% ice-cold methanol containing 0.1% formic acid, vortex-mixed, and centrifuged at 13,000 rpm for 10 min at 4°C. The clarified supernatant was then collected for metabolomic analysis.

Metabolomic profiling was performed using ultrafast liquid chromatography coupled to a TripleTOF 5600 high-resolution tandem mass spectrometer (AB SCIEX, UFLC-HRMS/MS), as previously described (**Deme et al., 2022**). Ten microliters of each sample were injected onto a Kinetex pentafluorophenyl (F5) column (Phenomenex, USA) and separated using a binary gradient mobile phase consisting of acetonitrile (eluent A) and deionized water (eluent B), both containing 0.1% formic acid. The gradient was programmed as follows: 0–3 min, 100% B; 3–13 min, 0–100% A; 13–19 min, 100% A; 19–19.1 min, return to 100% B; followed by a 4-min re-equilibration period.

Mass spectrometric data were acquired in both positive and negative electrospray ionization modes over a mass range of m/z 30–900 using information-dependent acquisition (IDA-HRMS/MS). Raw data were processed using SCIEX OS version 1.5 software integrated with the NIST 2017 Tandem Mass Spectral Library and the SCIEX Accurate Mass MS/MS Spectral Library 2.0 for peak detection, alignment, metabolite identification, and quantification. To establish a robust targeted metabolite reference library, pooled quality-control samples (n = 6) were analyzed to evaluate metabolite detectability, reproducibility, and analytical performance. Metabolites were included in the final targeted metabolite list only if they were detected in at least five of the six pooled runs and exhibited a coefficient of variation (CV) of less than 20% for peak area under the curve (AUC). These quality-control criteria ensured high analytical reliability, reproducibility, and quantitative precision. The finalized targeted metabolite list was then used as the reference library for metabolite identification and quantification in individual experimental samples. Missing values in experimental samples were imputed using the k-nearest neighbors (k-NN) algorithm. Internal standard performance was monitored throughout the study, and all internal standards demonstrated CVs below 20% across all analytical runs. Metabolite abundances were normalized to their corresponding internal standards prior to statistical analysis. In total, 173 metabolites were reliably quantified in each sample.

**mRNA Isolation, cDNA Library Preparation, and Bulk mRNA Sequencing**

Total mRNA was isolated from brain tissue using the RNeasy MicroKit (Qiagen, #74004) with qiazol-based extraction. Flash-frozen tissue was homogenized in 100 µL qiazol lysis reagent (Qiagen, #79306) with RNAprotect, incubated 5 minutes at room temperature, and 20 µL chloroform (Fisher, #C298-500) was added. After vigorous shaking for 20 seconds and a 3-minute room temperature incubation, samples were centrifuged at 4°C, 12,000 × g for 15 minutes. The aqueous phase (~60 µL) was recovered, combined with equal volume 70% ethanol, and transferred to a RNeasy MinElute column. Extraction proceeded according to manufacturer’s instructions, with steps conducted in a cold room unless otherwise specified.

RNA purity and concentration were assessed by NanoDrop spectrophotometry before submission to the Johns Hopkins Genetic Resources Core Facility for BioAnalyzer quality control, mRNA enrichment, cDNA library preparation, and sequencing. Only samples with RIN > 7 were used. mRNA was enriched using the NEBNext Poly(A) Magnetic Isolation Module (NEB #E7490L), cDNA libraries were prepared using the NEBNext Ultra II Directional RNA Library Prep Kit for Illumina (NEB #E7760), and sequencing was performed on an Illumina NovaSeq X Plus. Two technical replicates were generated per sample and aggregated during data analysis. Over 30,000 genes were quantified per sample.

**Data Processing**

Behavioral data were aggregated into 5-minute time bins using custom R code to permit area under the curve analyses. Latency data were defined as the first time a region was approached by the pup and were not binned.

Transcriptomic data were processed using a standard Trimgalore-HISAT2-Samtools-featureCounts pipeline^47–49^ to generate gene count matrices. Gene counts were analyzed via DESeq2^50^ for Differential gene expression analysis, control BA were used as reference unless otherwise noted, non-shrunken counts were used in pair-wise comparisons for initial identification of differentially expressed genes, and un-corrected p-values ares reported unless otherwise noted due to sample variation. For follow-up Weighted Gene Coexpression Network analysis (WGCNA), variance stabilized transformed gene counts were fed into the WGCNA package^51^package following previously established protocols. All sex-stratified analyses were conducted by subsetting DESeq2-normalized count matrices by sex prior to DGE and WGCNA.

Metabolomic data were log2 transformed then Pareto scaled before statistical analysis, metabolite set enrichment using the MetaboAnalystR package from Bioconductor, and Pearson correlation for metabolite-behavior correlation analysis.
Behavioral, metabolomic, and transcriptomic data were concatenated for machine learning-driven dimensionality reduction. All processing and quantification were conducted in R.

**Statistical Analysis**

For DAM duration and event data, pairwise comparisons across adversity conditions were conducted using the non-parametric Wilcoxon Rank Sum test, with non-parametric Welch’s ANOVA applied within each time point for body weight, maternal observation, MSA, and 2-CSP data. These analyses were conducted on pooled data, data stratified by sex separately (i.e., sex-specific), and across both sexes (i.e., sex-differences). Area under the curve was calculated for each binned behavioral measure and compared across conditions using the same approach. Because a subset of pups never approached a region of interest during the MSA test, latency data were analyzed as censored survival data using Kaplan-Meier estimation, with group differences assessed using the log-rank test and effect sizes estimated using Cox proportional hazards models with hazard ratios and 95% confidence intervals reported.

For SAM-LB body weight and behavior data, pairwise comparisons across drug and bedding conditions were conducted using the Wilcoxon Rank Sum test, with sex-stratified analyses conducted by subsetting within each sex prior to comparison. Metabolomic and transcriptomic data from the DAM samples were not corrected for multiple comparisons prior to pairwise comparisons by adversity treatment due to sample variation and experimental design complexity (adversity x sex x age). Additional analyses across and within sex were conducted within each developmental timepoint (infancy and adolescence) where appropriate. The specific statistical test applied depended on data distribution and whether comparisons were univariate or multivariate. Statistical significance was defined as p < 0.05 or p.adj < 0.05 for all analyses.
